# A baton-relay and proofreading mechanism for selective ER retrieval signal capture by the KDEL receptor

**DOI:** 10.1101/2021.03.21.436307

**Authors:** Andreas Gerondopoulos, Philipp Bräuer, Tomoaki Sobajima, Zhiyi Wu, Joanne L. Parker, Philip C. Biggin, Francis A. Barr, Simon Newstead

## Abstract

The KDEL-retrieval pathway captures escaped ER proteins with a KDEL or variant C-terminal signal at acidic pH in the Golgi and releases them at neutral pH in the ER. To address the mechanism of signal binding and the molecular basis for differences in signal affinity, we determined the HDEL and RDEL bound structures of the KDEL-receptor. Affinity differences are explained by interactions between the variable −4 position of the signal and W120, whereas initial capture of retrieval signals by their carboxyl-terminus is mediated by a baton-relay mechanism involving a series of conserved arginine residues in the receptor. This explains how the signal is first captured and then pulled into the binding cavity. During capture, retrieval signals undergo a selective proofreading step involving two gatekeeper residues D50 and E117 in the receptor. These mechanisms operate upstream of the pH-dependent closure of the receptor and explain the selectivity of the KDEL-retrieval pathway.

## INTRODUCTION

Stable maintenance of the luminal composition of the endoplasmic reticulum (ER) is crucial for the function of the secretory pathway (Ellgaard and Helenius, 2003). Because of the continuous flow of material from the ER to the Golgi, the essential chaperones and redox enzymes needed for protein folding in the ER lumen undergo dynamic retrieval from the Golgi apparatus (Gomez-Navarro and Miller, 2016). Conversely, secretory proteins destined for secretion and integral membrane proteins intended for other cellular compartments are not retained. This separation of secreted and retained cargo proteins involves signal-mediated sorting, whereby folded proteins destined for exit from the ER have active transport or exit signals, and proteins to be retained in the ER have signals for retrieval (Barlowe, 2003; Gomez-Navarro and Miller, 2016). For membrane proteins, cytoplasmic signals can directly engage with the selective vesicle coat complexes required for transport between the ER and Golgi. For luminal proteins, this information has to be relayed by a transmembrane receptor that serves as an intermediary to the cytoplasmic coat protein complexes (Dancourt and Barlowe, 2010). In the archetypal KDEL-retrieval system, a 7-transmembrane receptor captures escaped ER luminal proteins carrying a C-terminal KDEL or variant tetrapeptide sequence in the acidic pH of the Golgi (Munro and Pelham, 1987; Semenza et al., 1990). Signal binding to a luminal cavity in the receptor triggers a conformational change in its cytoplasmic face that exposes a lysine motif recognised by the COP I coat complex (Bräuer et al., 2019). Release of the signal in the neutral pH environment of the ER results in a reversal of this conformational change, burying the lysine motif, and exposing a patch of aspartate and glutamate residues presumed to form a COPII-binding ER exit signal for the receptor (Bräuer et al., 2019; Newstead and Barr, 2020). Hence, the KDEL receptor cycles between the ER and Golgi capturing escaped ER proteins in a dynamic retrieval process (Dean and Pelham, 1990; Lewis and Pelham, 1992; Townsley et al., 1993; Zagouras and Rose, 1989). The rapid recycling of the receptor means it does not need to be stoichiometric with the ER concentration of retained proteins, only present at levels sufficient to capture escaped proteins that reach the Golgi (Newstead and Barr, 2020). However, ER resident proteins differ widely in abundance, yet, remarkably, this does not pose a problem for efficient retention of low abundance proteins. One possible explanation for this is the presence of HDEL and RDEL variants of the canonical KDEL signal with different binding affinities (Scheel and Pelham, 1998; Wilson et al., 1993). However, despite extensive mutation and structural analysis the molecular basis and functional significance of these affinity differences remains unclear (Bräuer et al., 2019; Townsley et al., 1993). Complicating this picture, in some organisms including the yeasts *Kluyveromyces lactis* and *Schizosaccharomyces pombe*, DDEL and ADEL variants are used as ER retrieval signals (Pidoux and Armstrong, 1992; Semenza and Pelham, 1992). Comparative analysis of the budding yeast *Saccharomyces cerevisiae* HDEL- and *K. lactis* DDEL-receptors implicated a luminal region including a key variant residue, D50 in the human receptor, in selectivity for DDEL (Lewis et al., 1990; Semenza et al., 1990; Semenza and Pelham, 1992). Mutation of D50 to cysteine in the human receptor resulted in reduced binding affinity for KDEL, RDEL and HDEL (Scheel and Pelham, 1998). However, recent structure determination of the chicken receptor with a bound TAEKDEL peptide indicates this residue sits on the luminal surface of the receptor and does not make contact with any portion of the signal (Bräuer et al., 2019). Thus, although it is clear that the specificity of ER retrieval is encoded by the KDEL receptor, the molecular basis for the recognition of different signal variants remains unclear.

Our previous work has shown the KDEL receptor has a transporter-like architecture and undergoes pH-dependent closure around cognate retrieval signals (Bräuer et al., 2019; Newstead and Barr, 2020). However, the molecular basis for affinity differences for retrieval signal variants and any functional significance these differences may create, was not explained by that work or other previous studies. Furthermore, how signals are initially captured and selected from other sequences remains enigmatic. To answer these related questions, we solved structures of the human KDEL receptor in complex with both HDEL and RDEL retrieval signals, and performed a combination of computational and cell biological analysis. Based on this data, we can break down the retrieval signal recognition process into a series of steps for signal proofreading and initial capture of the free carboxyl terminus, followed by full engagement with the binding cavity and finally pH-dependent closure and activation of the receptor.

## RESULTS

### ER retrieval signals in mammalian cells

To understand how the KDEL-receptor differentiates between cargo proteins, we first sought to define the major signal variants used in mammalian cells. For this purpose, we exploited luminal ER proteome datasets to investigate the relative abundance of retrieval signal variants (Itzhak et al., 2017; Itzhak et al., 2016). This confirmed that KDEL, HDEL and RDEL are the major variants in mammals, and the frequency of ER resident proteins with these variants of the retrieval signal at the −4 position is approximately equal (Figure 1a). However, this does not reflect the abundance of the proteins carrying the signal. Strikingly, the total concentration of KDEL bearing proteins is over five-fold higher than either HDEL or RDEL (Figure 1b). This largely reflects a small number of highly abundant ER-resident chaperones, BIP, PDI and calreticulin (Figure 1 – supplement 1a and b). Each of these proteins is present in the 5-10 µM range, far more abundant than the dominant KDELR receptor 2 (KDELR2) species which is estimated to be 0.2-0.3 µM (Figure 1 – supplement 1c). In total, the concentration of retrieval signals thus exceeds that of the receptor by at least two orders of magnitude. In good agreement with previously reported studies on the mammalian KDEL receptor (Scheel and Pelham, 1998; Wilson et al., 1993), we found that HDEL has the highest affinity for the receptor K_D_ 0.24 µM, followed by KDEL K_D_ 1.94 µM and RDEL K_D_ 2.71 µM (Figure 1c). Previous work has suggested DDEL binds to semi-purified human KDEL receptors in membrane fractions and can function as a retrieval signal when the receptor is overexpressed at high level in COS7 cells (Lewis and Pelham, 1992; Wilson et al., 1993). However, we find that DDEL binds with 60-fold lower affinity than HDEL (K_D_ 14.9 µM) (Figure 1c), in agreement with other data for purified KDEL receptors (Scheel and Pelham, 1998). Thus, the receptor binds to the HDEL sequence with one order of magnitude greater affinity than the canonical KDEL ligand present on the most abundant ER resident proteins. Despite this difference in affinities, mScarlet fusions with KDEL, RDEL or HDEL signals all triggered similar changes to the steady-state distribution of the KDEL receptor in cells, driving almost complete retrieval from the Golgi to the ER (Figure 1d and 1e). By contrast, expression of ADEL or DDEL had little effect on the Golgi-ER distribution of the receptor (Figure 1d and 1e). In line with these effects on the receptor, the mScarlet-KDEL, RDEL and HDEL ligands were retrieved to the ER, whereas ADEL and DDEL showed predominantly Golgi and punctate localisation consistent with secretion (Figure 1d). These latter observations explain why there are no verified examples of endogenous proteins using ADEL and DDEL retrieval signals in mammalian cells.

**Figure 1.**
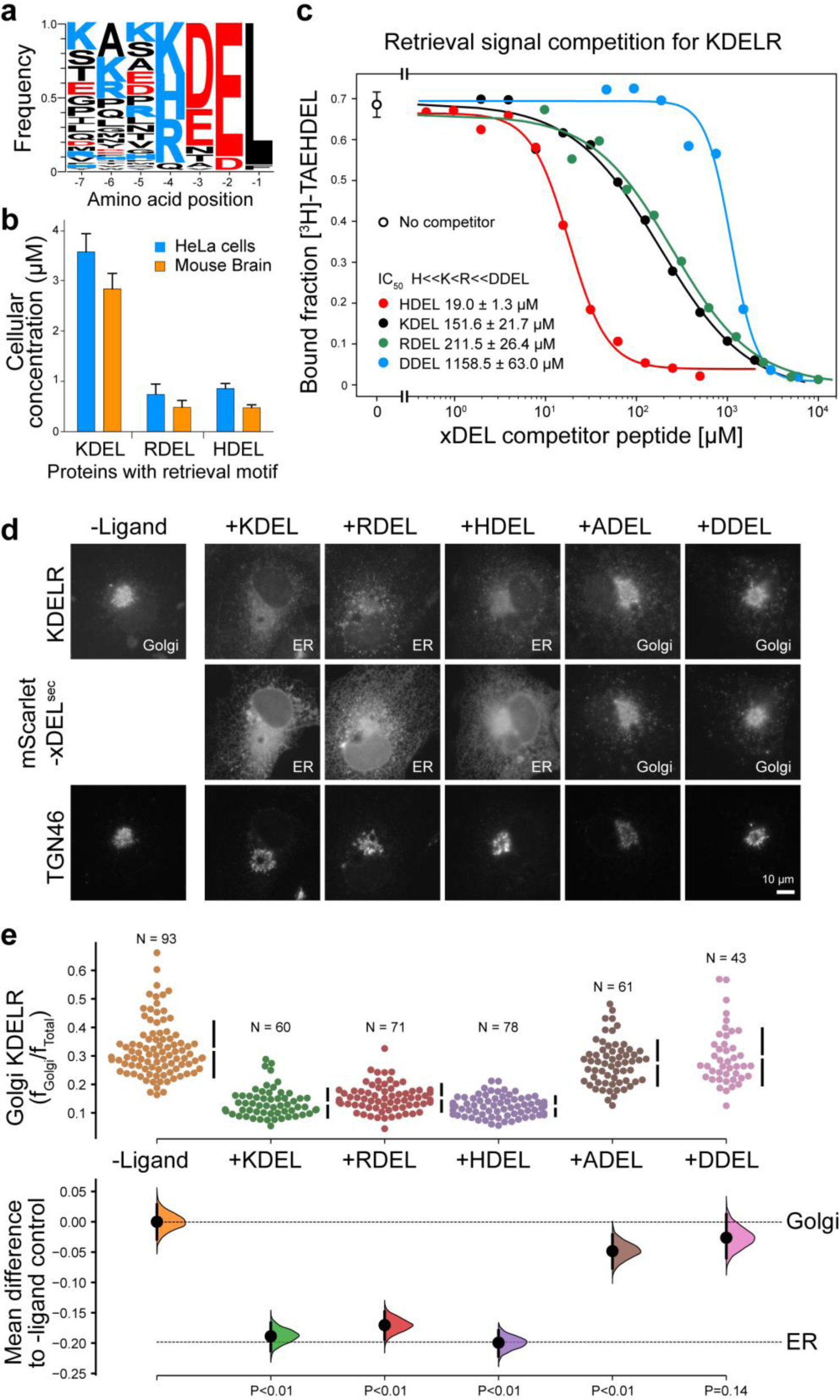
ER retrieval signal abundance and affinity are not correlated. **a.** Sequence logos for ER resident proteins with C-terminal KDEL retrieval signals and variants thereof calculated using frequency or protein abundance (Itzhak et al., 2017; Itzhak et al., 2016). **b.** Combined cellular concentrations of ER resident proteins with canonical KDEL, RDEL and HDEL retrieval sequences in HeLa cells and mouse brain. **c.** Competition binding assays for [^3^H]-TAEHDEL and unlabelled TAEKDEL, TAERDEL and TAEHDEL to the KDEL receptor showing IC_50_ values for the competing peptides. These were used to calculate apparent K_D_ using the Cheng-Prusoff equation (Cheng and Prusoff, 1973). **d.** Endogenous KDEL receptor redistribution was measured in the absence (-ligand) or presence of K/R/H/A/DDEL (mScarlet-xDEL^sec^). TGN46 was used as a Golgi marker. Scale bar is 10µm. **e.** The mean difference for K/R/H/A/DDEL comparisons against the shared no ligand control are shown as Cummings estimation plots. The raw data for the fraction of KDEL receptor fluorescence in the Golgi is plotted on the upper axes with sample sizes and p values.

Given its higher affinity, why then is HDEL not the dominant ER retrieval signal, especially for crucial ER proteins such as BIP, PDI and calreticulin? We tested the idea that due to its higher binding affinity, increasing the concentration of HDEL bearing proteins would effectively compete for KDEL receptors in the Golgi, and prevent efficient ER retrieval of KDEL and RDEL containing proteins. To do this we used our series of variant xDEL signals, where x at the −4 position is either K, R, H, A or D. When expressed in cells, KDEL, RDEL and HDEL are retained in the cell, whereas ADEL and DDEL are mostly secreted (Figure 1 – supplement 1d and e). With the exception of HDEL this is broadly in line with their respective binding affinities. Despite binding to the receptor with a higher affinity (Figure 1c), HDEL was less efficiently retained than either KDEL or RDEL (Figure 1 – supplement 1d and e). We then examined the effect of these ligands on the major ER proteins BIP and PDI as well as the less abundant chaperones ERP72 and ERP44 ((Figure 1 – supplement 1a). As predicted, ADEL and DDEL had little effect on ER retention, while HDEL caused secretion of all four proteins (Figure 1 – supplement 1f).

These results indicate that the retrieval system is selective yet not optimised for binding affinity, and instead has evolved to ensure optimal retrieval of a broad cohort of proteins of widely differing abundance. In human cells, ADEL and DDEL do not bind to the receptor with high affinity and do not function as retrieval signals, suggesting specific recognition of the −4 position is a key determinant for binding. Previously, it has been suggested that complementary charges at receptor position 50 and the −4 position of the signal explain this specificity (Lewis and Pelham, 1992; Semenza and Pelham, 1992). However, this mechanism does not obviously explain how ADEL, with no charged residue at the −4 position, functions as a signal in some organisms. How signal selectivity is achieved is therefore a crucial question we need to answer.

### HDEL and RDEL signals bind similarly to the canonical KDEL variant

To understand the molecular basis for the affinity differences between retrieval signal variants, we determined structures for the chicken KDELR2 bound to HDEL and RDEL signals. These structures with TAEHDEL and TAERDEL peptides have resolutions of 2.24 and 2.31 Å, respectively (Figure 2a-2c and Table S1). In both instances the overall structure of the receptor is similar to our previous complex with the TAEKDEL peptide (Figure 2d), with a root mean square deviation (R.M.S.D.) of 0.223 and 0.153 Å over 200 C_α_ atoms for the HDEL and RDEL structures, respectively. Both HDEL and RDEL peptides are bound in a vertical orientation with respect to the membrane, with the side chains clearly resolved in the electron density map (Figure 2 – supplement 1a and b). Both the HDEL and RDEL peptides interact with the receptor through the same salt bridge interactions seen for the KDEL peptide (Figure 2b-2d). Superimposing the three peptides reveals little movement of the peptide at the −1 and −2 positions when bound to the receptor (Figure 2e). For RDEL, we observe slight movement of the backbone C_α_ atom of the peptide to accommodate the larger arginine side chain, resulting in a minor repositioning of the glutamate at the −3 position in the receptor. Nonetheless, the position of the positive charge at the −4 position on all three peptides is identical relative to E117 and W120 within the receptor, supporting the view that a salt bridge is formed with E117 on TM5. D50 previously proposed to be important for recognition of the −4 position is at >5 Å distance, outside the region depicted in the figures, indicating it is unlikely to form a salt bridge and directly contribute to binding of the retrieval signal. Some studies have suggested the core tetrapeptide retrieval motif should be extended to include the −5 and −6 positions (Alanen et al., 2011). However, these positions are not conserved in retained ER luminal proteins (Figure 1a). In our structures, the glutamate at the −5 position sits close to S54, but would not obviously increase the binding affinity, whereas no contacts are made to the −6 position. In all cases, the −1 position leucine residue and free carboxy terminus form interactions to R47 and Y48 on TM2, as well as R159 and Y162 on TM6. The glutamate at position −2 forms a further salt bridge interaction to R5 on TM1 and a hydrogen bond to W166 on TM6, whereas the aspartate at −3 forms a salt bridge with R169, also on TM6. For the histidine side chain at the −4 position of HDEL, the imidazole group is predicted to form a π-π stacking interaction with W120 (Figure 2b). In comparison, the RDEL arginine side chain sits in the same position as the amine group of the KDEL sequence and could thus interact with W120 via a cation-π interaction and E117 via a classical salt bridge interaction (Figure 2c). We therefore conclude that both E117 and W120 play a role in retrieval signal binding, and the only major difference between the HDEL, RDEL and KDEL signals is the precise nature of the interaction with W120 indicating that this may be a critical residue to explain the affinity of different signals.

**Figure 2.**
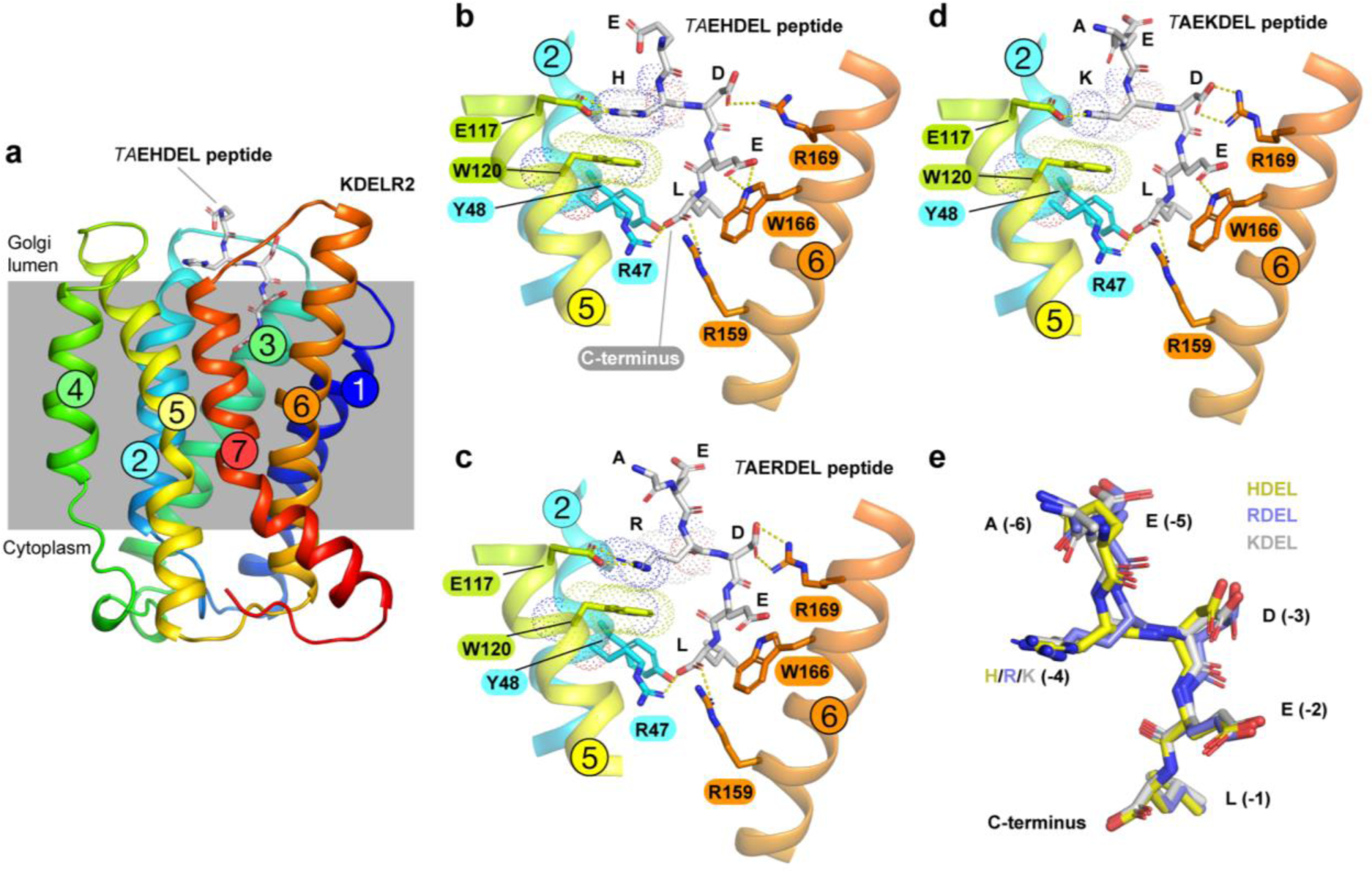
Structures of the KDEL receptor bound to HDEL and RDEL retrieval signals. **a.** Crystal structure of chicken KDELR2 viewed from the side with the transmembrane helices numbered and coloured from N-terminus (blue) to C-terminus (red). The predicted membrane-embedded region of the receptor is indicated by a grey shaded box, with labels at the lumenal and cytoplasmic faces. The TAEHDEL peptide is shown in stick format, coloured grey. **b.** Close up views of bound TAEHDEL (this study), **c.** TAERDEL (this study) and **d.** TAEKDEL (PDB:6I6H) peptides bound to the receptor are shown with contributing side chains labelled. Hydrogen bonds are indicated as dashed lines. The molecular orbitals of W120 and the −4 histidine on the peptide are shown as a dotted surface. **e.** Superposition of the HDEL, RDEL and KDEL peptides reveals near identical binding position within the receptor. Retrieval signal side chains are numbered counting down from the C-terminus.

### Probing the importance of E117 and W120 for signal binding

To directly test the requirement for E117 and W120 in signal recognition, ligand binding assays using recombinant chicken wild type, E117 or W120 mutant KDELR2 were performed (Bräuer et al., 2019). All proteins had similar thermal stability indicating they were correctly folded. For the wild type receptor at pH 5.4, K_D_ for KDEL and HDEL signals were 1.9 ± 0.46 µM and 0.26 ± 0.04 µM, respectively (Figure 3a and 3b). Conservative substitution of E117 with aspartate resulted in a slight reduction in binding for both KDEL and HDEL, K_D_ of 2.1 ± 0.33 µM and 0.52 ± 0.02 µM, respectively (Figure 3a and 3b). Substitution of E117 with alanine had a greater effect on KDEL binding, K_D_ ∼9.3 ± 1.0 µM, compared to HDEL, K_D_ 0.52 ± 0.02 µM (Figure 3a and 3b). This suggested that any salt bridge to E117 plays a greater role for KDEL than HDEL signals.

**Figure 3.**
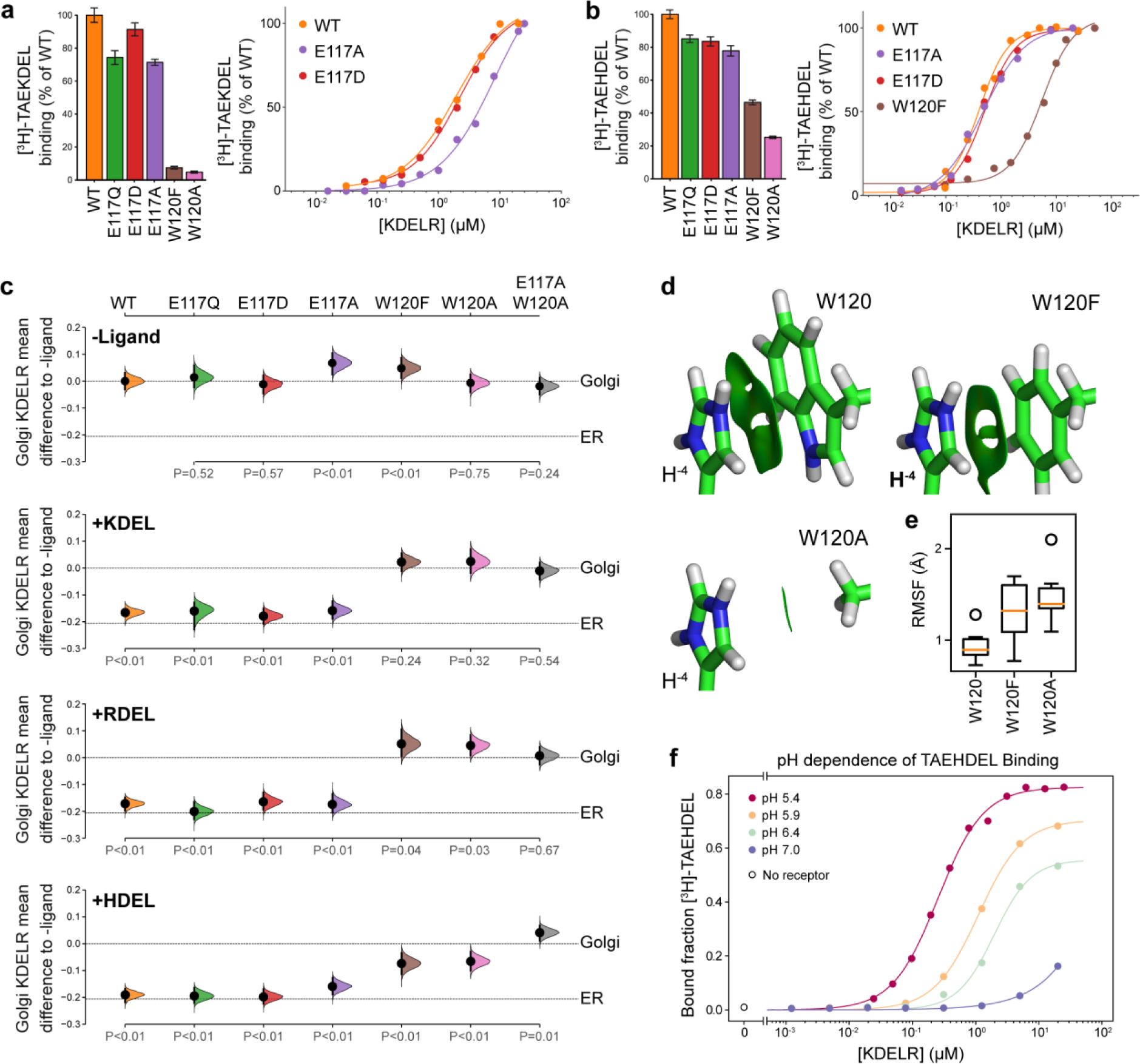
Roles of KDEL receptor E117 and W120 in retrieval signal binding and function in cells. **a.** Normalised binding of [^3^H]-TAEKDEL and **b.** [^3^H]-TAEHDEL signals to purified WT and the indicated E117 and W120 mutant variants of chicken KDELR2. Bar graphs show mean binding ± SEM (n=3). Line graphs show titration binding assays. **c.** The fraction of WT, E117 and W120 mutant KDEL receptor localised to the Golgi was measured before (no ligand) and after challenge with different retrieval signals (K/R/HDEL) as indicated. Effect sizes are shown as the mean difference for K/R/HDEL comparisons against the shared -ligand control with sample sizes and p-values. **d.** The π-π interactions between W120 and the histidine were visualised using reduced density gradient analysis. The wild-type W120 exhibit stronger π-π interactions compared with W120F, while W120A shows no π-π interactions. **e.** When W120 is changed to phenylalanine, the protonated histidine has a higher root mean squared fluctuation (RMSF) in the binding pocket, which is further increased for the W120A substitution. **f.** Binding of [^3^H]-TAEHDEL to the KDEL receptor was measured at pH5.4-7.0 and is plotted as a function of receptor concentration.

We next examined the contribution of W120 to signal recognition. Tryptophan side chains have long been recognized as important contributors in protein ligand interactions, as they are capable of interacting with ligands via both aromatic and charged forces (Dougherty, 1996; Liao et al., 2013; Okada et al., 2001). Our structures show that the −4 position histidine, arginine or lysine side chain of the human retrieval signal variants can in principle interact favourably with W120 via cation-π interactions. We also reasoned that given the additional π-π stacking observed with the imidazole group in the crystal structure, this interaction might explain the increased affinity observed for the HDEL signal variant. Mutation of W120 to alanine resulted in loss of binding to the KDEL peptide and it was not possible to calculate a K_D_ (Figure 3a). For the HDEL peptide, binding was reduced to 20% confirming that W120 plays an important role in mediating receptor-peptide interactions (Figure 3b). Consistent with the hypothesis that the −4 histidine of HDEL undergoes π-π stacking interactions with W120, conserved substitution to phenylalanine supported 50% HDEL binding with K_D_ 5.5 ± 0.57 µM, whereas no interaction was observed with the KDEL peptide (Figure 3a and 3b). Thus, W120 plays a crucial role in binding of both KDEL and HDEL signals and may explain the higher affinity of the receptor for HDEL. By contrast, E117 is less important and it is unclear why it is a conserved feature of the binding site.

To analyse whether the properties measured using purified components *in vitro* reflect the behaviour of the KDEL receptor and retrieval system *in vivo*, we analysed the ability of these same variants in the human KDEL receptor to differentiate between human retrieval signal sequences in a cellular ER retrieval assay. All the receptor mutants tested reached the Golgi apparatus supporting the view they are able to fold and exit the ER (Figure 3c, -Ligand, and Figure 3 – supplement 1a). The WT receptor showed robust retrieval to the ER in response to model cargo proteins bearing KDEL, RDEL or HDEL retrieval sequences (Figure 3c and Figure 3 – supplement 1a-d). Receptors with conservative (E117Q and E117D) or non-conservative (E117A) substitutions at E117 were efficiently retrieved to the ER with KDEL, RDEL or HDEL signal variants (Figure 3c and Figure 3 – supplement 1b-d). By contrast, receptors with mutations at W120A and W120F did not respond to KDEL and RDEL signals and showed greatly reduced response to HDEL (Figure 3c and Figure 3 – supplement 1b-d). The residual response to HDEL was abrogated in a double E117A/W120A mutant receptor (Figure 3c and Figure 3 – supplement 1b-d). This *in vivo* behaviour is in good agreement with the changes to affinity measured using *in vitro* binding assays (Figure 3a and 3b), and supports the view that W120 is of greater importance for ligand binding and ER retrieval.

To provide further support for this conclusion, we investigated the free energy of interaction between the histidine side chain of the retrieval signal and W120 of the receptor. Protonation of the HDEL histidine is a crucial consideration since retrieval signal binding to the receptor occurs at acidic pH in the Golgi. We therefore asked if the protonation state of the histidine is important for binding affinity. Molecular mechanics-based alchemical transformation was used to compute the free energy difference of changing the lysine in KDEL to different protonation states of the histidine in HDEL. The binding free energy of HDEL is −1.8 ± 1.4 kcal.mol^-1^ stronger than the KDEL signal (Table S2), which is in good agreement with the expected −1.3 kcal/mol free energy difference derived from measured K_D_ values for KDEL and HDEL. The preference for HDEL of −1.9 ± 0.2 kcal.mol^-1^ is mainly attributed to the protonated histidine which makes favourable cation-π interactions with W120 (Table S2, HIP). In agreement with the experimental data (Figure 3b and 3c), the W120F mutation, which is anticipated to preserve the cation-π interactions, reduced but did not abolish the preference for HDEL to −0.7 ± 1.6 kcal.mol^-1^, notwithstanding the large error on this calculation. Furthermore, the W120A mutation which eliminates the cation-π interactions, greatly reduced the preference for HDEL to −0.3 ± 0.9 kcal.mol^-1^.

To quantify the strength of the π-π and cation-π interactions between W120 variants and the histidine, we decomposed the interactions using symmetry-adapted perturbation theory from quantum mechanics. Although both W120 and W120F form π-π and cation-π interactions with protonated histidine, W120F exhibits ∼1.5 kcal/mol weaker π-π interactions and ∼0.5 kcal/mol weaker cation-π interactions with the histidine (Figure 3d and Table S3). The consequence of these changes is that for W120F higher root mean squared fluctuations are seen (Figure 3e), indicative of less rigid binding. These fluctuations are further increased for W120A (Figure 3e), consistent with its greater effect on signal binding. These results support the hypothesis that the π-π interactions between the protonated histidine sidechain and W120 explain the higher affinity observed for HDEL signals. Further support for this interpretation comes from *in vitro* analysis of the pH-dependence of HDEL binding. At pH6.4 HDEL shows ∼60% maximal binding to the receptor (Figure 3f), compared to <20% seen at the same pH for KDEL (Bräuer et al., 2019). The level of HDEL binding seen at pH 7 would saturate the KDELR receptor in the ER if the most abundant luminal proteins such as BIP carried this signal variant. Our observation that W120 is also necessary for recognition of KDEL indicates that cation-π interactions to W120, rather than a salt bridge to E117, is the crucial determinant for recognition of the −4 position.

### E117 plays a role in KDEL receptor selectivity

This mode of signal binding involving W120 is different than previously proposed, where charge complementarity between D50 in the receptor and the −4 position of the signal was thought to be a key determinant of specificity in ER retrieval (Lewis and Pelham, 1992; Scheel and Pelham, 1998; Semenza and Pelham, 1992). However, as our crystal structures show, D50 is outside the immediate binding region for all retrieval signal variants and therefore unlikely to directly contribute to binding. Thus, the precise roles of D50 and E117 remain enigmatic. In this regard the behaviour of ADEL signals is noteworthy due to the simple methyl side chain. Comparison of different retrieval signals shows that ADEL does not activate the wild type human KDEL receptor (Figure 1d and 1e). The simplest explanation for this finding is that the −4 position is crucial for high affinity binding of retrieval signals to the human receptor. However, this view is unlikely to be correct. First, the KDEL, RDEL and HDEL bound receptor structures do not support the view that recognition of the −4 position requires D50, and instead provide an alternative possibility where E117 fulfils this role. However, our biochemical and functional data show that E117 does not contribute greatly to signal binding affinity or retrieval in cells (Figure 3a-3c). Therefore, rather than selecting for the sequence, E117 may be more important to select against unwanted signal variants. To test this idea, we examined the response of E117A mutant receptors to variant ADEL and DDEL signals. Remarkably, the E117A mutant receptor relocated to the ER in response to both KDEL and ADEL, but not DDEL signals (Figure 4a and 4b). In *S. pombe* and *K. lactis*, organisms where ADEL and DDEL are used for ER retrieval, the E117 position is either an asparagine or a glutamine residue, and we therefore tested E117N and E117Q mutants next. Similar to the results with E117A, E117Q and E117N receptors move to the ER in response to KDEL or ADEL signals, yet interestingly still failed to respond to DDEL (Figure 4a and 4b). Ligand expression was in a similar range in all instances (Figure 4 – supplement 1), and in the absence of ligand all three mutant receptors localised to the Golgi with a low ER background indicating normal folding and ER exit (Figure 4a).

**Figure 4.**
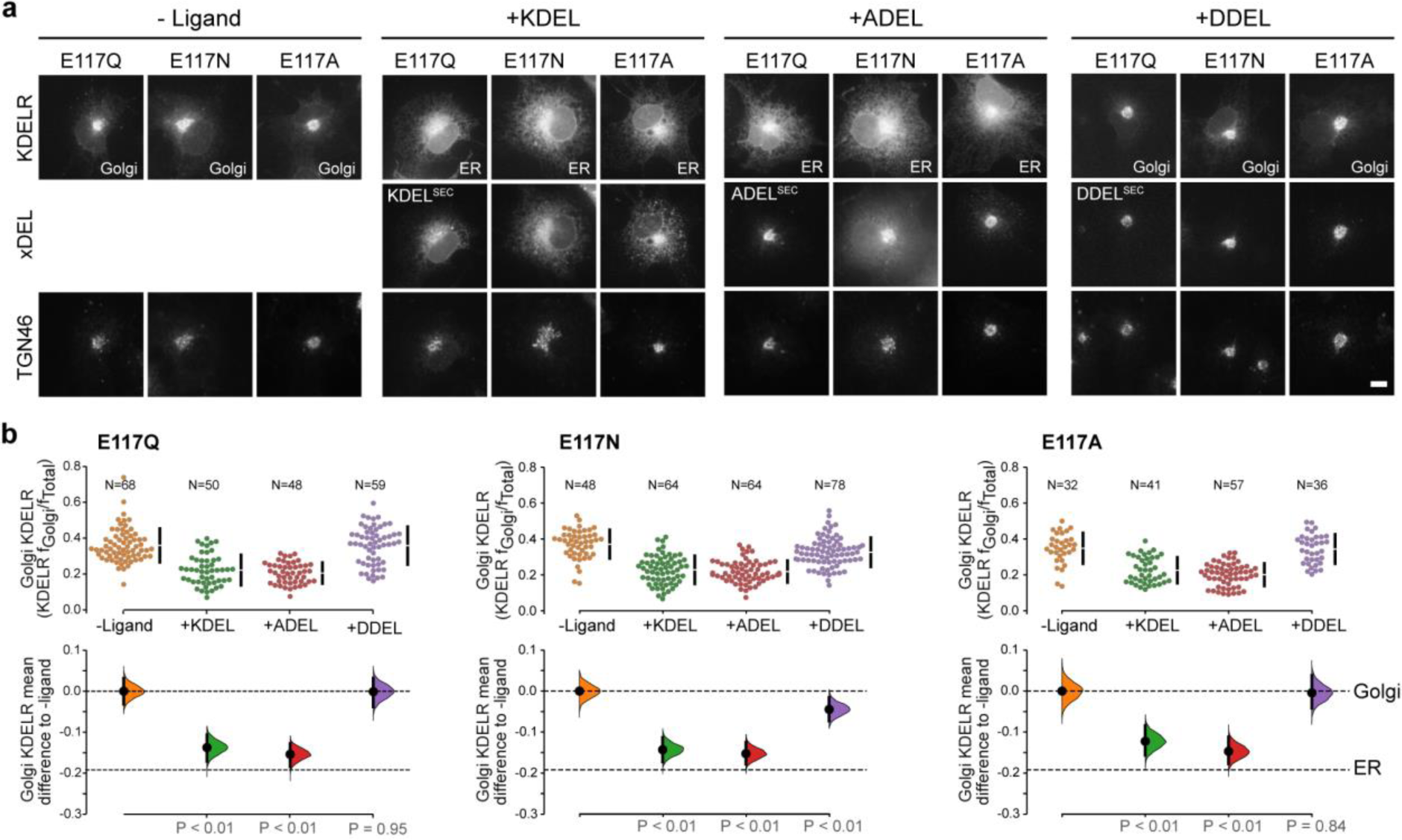
KDEL receptor E117 mutants show reduced selectivity for retrieval signals. **a.** E117Q, E117N or E117A mutant KDEL receptors were tested for K/A/DDEL-induced redistribution from Golgi to ER. KDEL receptor distribution was followed in the absence (-ligand) or presence of K/A/DDEL^sec^. TGN46 was used as a Golgi marker. Scale bar is 10µm. **b.** The fraction of E117Q, E117N or E117A mutant KDEL receptor localised to the Golgi was measured before (no ligand) and after challenge with different retrieval signals (K/A/DDEL). Effect sizes are shown as the mean difference for K/A/DDEL comparisons against the shared -ligand control with sample sizes and p values.

Thus, E117 is important for determining which signals are rejected by the wild type human receptor based on the −4 position of the signal, but does not appear to play a major role in binding affinity. ADEL must bind to the E117A mutant receptors via the “DEL” tri-carboxylate portion of the retrieval signal, suggesting this region may be the major contributor to binding affinity for all signal variants. For HDEL, the protonated histidine side chain makes additional π-π interactions with W120 to bind with higher affinity. Importantly, the lack of response to DDEL shows that signal selection and recognition must involve additional features in the *S. pombe* and *K. lactis* receptor, and we investigated this question further.

### A charge screening mechanism for signal differentiation by the KDEL receptor

To identify additional features that might play a role in signal selection, we performed a comparison of the receptors and most abundant cognate ligands of the HSPA5/BIP family of ER resident proteins in different species. Although most regions of the receptor are highly conserved, as noted previously (Semenza and Pelham, 1992), sequence alignment reveals two regions where there is covariation that may be related to the cognate tetrapeptide retrieval signal (Figure 5a). In receptors recognising ADEL and DDEL, D50 is changed for asparagine, E117 for glutamine or asparagine, and position 54 is a positively charged arginine or lysine rather than a polar side chain (Figure 5a). To understand the consequences of these changes we examined their positions relative to the bound TAEHDEL signal (Figure 5b). This reveals that E117 and S54 sit close to the −4 histidine and −5 glutamate, respectively and D50 is over 5 Å away from any residue in the signal in the final bound state (Figure 5b). Analysis of the charge distribution across the surface of the receptor shows a negatively charged feature above the positively charged binding cavity occupied by the DEL portion of the signal, with the −4 residue sited at the boundary to these two regions (Figure 5c). Strikingly, progressive introduction of changes in the human receptor to mimic the *K. lactis* receptor, D50N S/N54K E117Q erodes the negatively charged luminal feature (Figure 5d).

**Figure 5.**
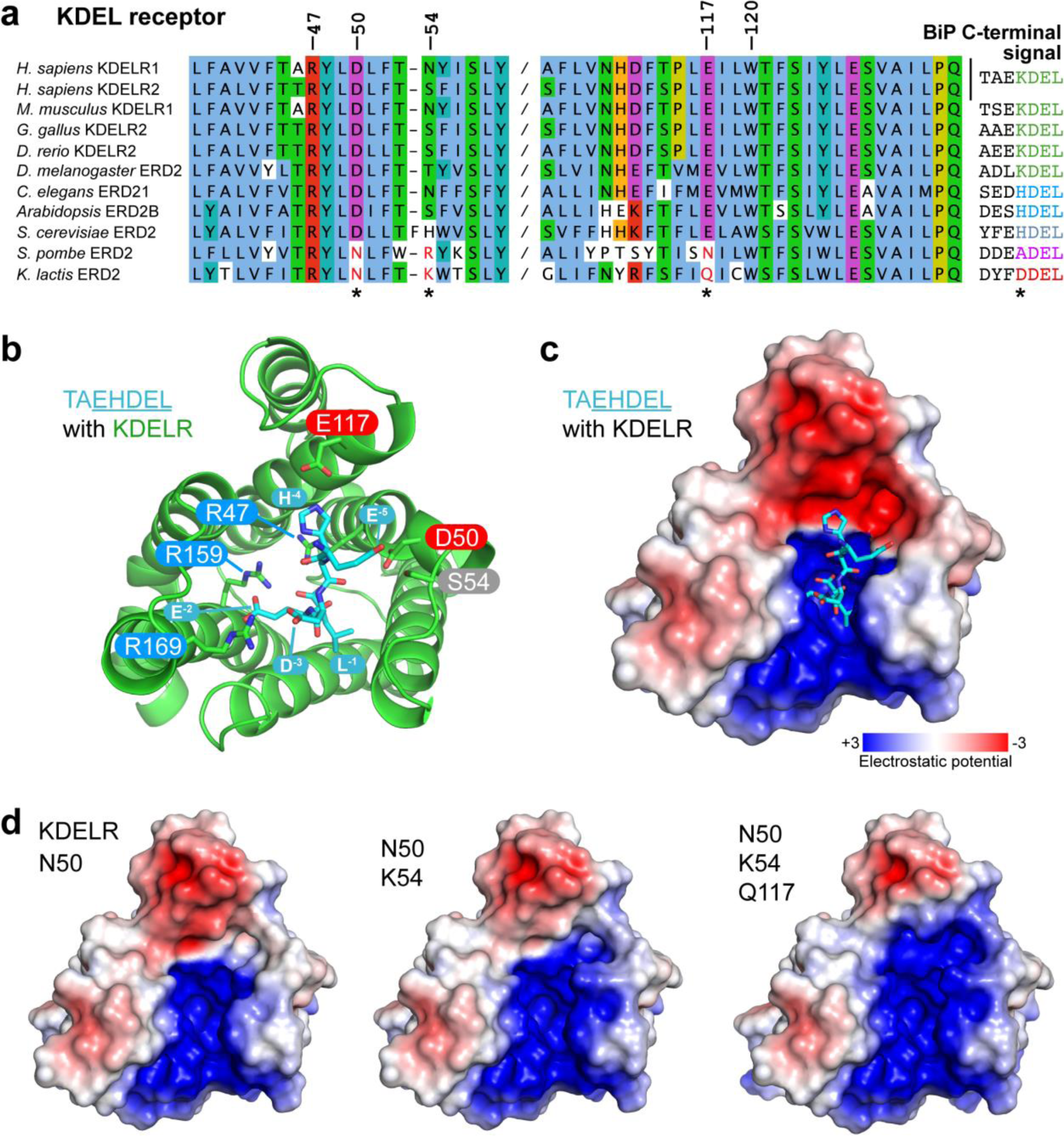
Charge distribution across the luminal opening of the KDEL receptor binding pocket. **a**. KDEL receptor sequence alignment showing two regions centred around amino acid D50 and W120 of the human proteins. Cognate retrieval signal variants are shown to the right of the alignment. **b.** The structure of the KDEL receptor with bound TAEHDEL highlighting key residues involved in ligand binding and variant residues D50, N54 and E117. **c.** The charged surface for the WT KDEL receptor and **d.** N50, N50/K54 and N50/K54/Q117 mutants is shown.

One simple explanation for this feature is that it extends the binding site to impart specificity for the region upstream of the core KDEL signal. However, analysis of different classes of ER luminal proteins from yeast and animal cells does not provide strong support for this possibility. The upstream sequences of many abundant ER proteins including human and yeast HSPA5/BIP homologues are acidic in nature, and not basic (Figure 5a and Figure 5 – supplement 1a), making any interaction unfavourable. For the human signal, the −4 position is crucial and mutation to A or D abolishes ER retrieval of the receptor (Figure S5b and Figure 5 – supplement 1d). Conversely, *S. pombe* and *K. lactis* BIP ADEL and DDEL signals become functional with the human receptor if the −4 position is changed to lysine confirming this is the critical residue, independent of upstream sequences (Figure 5 – supplement 1c and d). In *K. lactis* BIP the −5 position is a bulky aromatic residue rather than a charged residue. Previous work has suggested that the budding yeast FEHDEL signal with a bulky aromatic residue at the −6 position does not function in mammalian cells (Wilson et al., 1993), however consistent with our other data we find this HDEL variant is also functional (Figure 5 – supplement 1c). Extending this analysis to human FKBP family proteins with even more diverse upstream sequences reveals no obvious pattern of conservation other than the canonical C-terminal HDEL or HEEL retrieval signal (Figure 5 – supplement 1e).

To directly test the role of the charged luminal surface in signal selection, we made a series of mutants introducing the changes seen in *K. lactis* and *S. Pombe* into the human receptor and tested these against KDEL, ADEL and DDEL signals. A single D50N mutation abolished the response to all signal variants and the receptor remained in the Golgi (Figure 6a and 6b). Thus, like E117, D50 is not the sole determinant of signal selectivity. Similarly, N54K reduced the response to KDEL but did not result in ADEL or DDEL recognition (Figure 6a and 6b). A D50N N54K double mutant showed a loss of specificity and gave an intermediate response to KDEL, ADEL and DDEL signals, showing that it is possible to uncouple binding from selectivity at the −4 position. We then combined D50N or N54K with E117Q mutations. These double mutant receptors showed switched specificity towards ADEL and DDEL with only a residual response to KDEL (Figure 6a and 6b). Combination of D50N N54K and E117Q improved the response to ADEL and DDEL and further reduced that towards KDEL (Figure 6a and 6b). Comparable results were obtained with a *S. pombe* like D50N N54R E117N triple mutant receptor (Figure 6 – supplement 1a and b). Both these altered specificity receptors responded to the cognate ADEL or DDEL variant of BIP for that organism, a response that was abolished solely by mutation of the −4 position of the signal (Figure 6 – supplement 1a and b).

**Figure 6.**
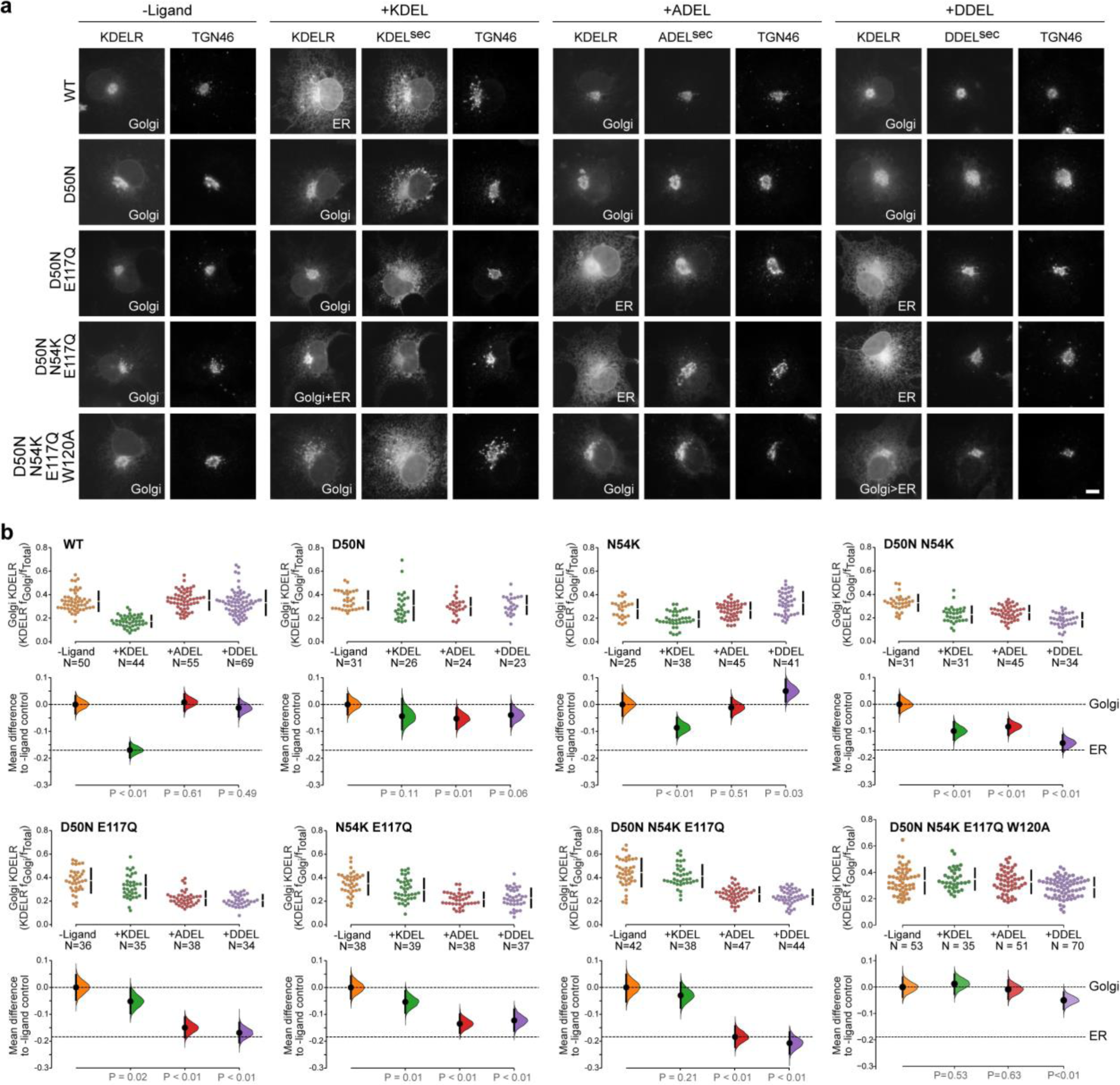
Re-engineering the selectivity of the human KDEL receptor for ADEL and DDEL signals. **a.** WT and a series of *“K. lacti*s”-like mutant KDEL receptors were tested for K/A/DDEL-induced redistribution from Golgi to ER. KDEL receptor distribution was followed in the absence (-ligand) or presence of K/A/DDEL^sec^. TGN46 was used as a Golgi marker. Scale bar is 10µm. **b.** The fraction of WT and mutant KDEL receptor localised to the Golgi was measured before (no ligand) after challenge with different retrieval signals (K/A/DDEL). Effect sizes are shown as the mean difference for K/A/DDEL comparisons against the shared -ligand control with sample sizes and p values.

These results indicate that the −4 position of the signal is read out during initial signal binding and is important for exclusion of unwanted signals, but is less important for binding affinity. We therefore tested whether the mode of ADEL and DDEL binding to the switched specificity receptors still involves W120. A D50N N54K E117Q W120A mutant receptor does not relocate from the Golgi to the ER with KDEL and ADEL signals and shows only a small response to the DDEL signal (Figure 6a and 6b). Together, these findings suggest a common mode of binding for all retrieval signal variants through residues conserved in all species. Specificity for the −4 position is largely achieved through a proofreading mechanism involving two gatekeeper residues, D50 and E117, as the signal enters the ligand binding cavity. An E117A substitution partially uncouples this mechanism and allows ADEL binding, whereas both D50 and E117 residues have to be changed to allow DDEL binding. Bringing together all our observations to this point, we conclude that the luminal surface of the receptor plays a crucial role in signal selectivity prior to adoption of the final activated state, perhaps by determining the rate of signal association from solution.

### Initial retrieval signal capture by the free carboxyl terminus

To explore the initial interaction of retrieval signals with the KDEL receptor we simulated an all-atom model of the KDEL signal with a free C-terminal carboxylate engaging with the receptor (Supplemental movie 1). This simulation shows that the signal initially encounters the receptor through a salt bridge interaction from its C-terminal carboxyl group with R169 on TM6 of the receptor (Figure 7a, i.). The C-terminal carboxyl group then moves to engage R5 (Figure 7a, ii.), shortly followed by interaction of the −2 glutamate with R169 (Figure 7a, iii.). Finally, the C-terminus engages with R47 on TM4 enabling the −3 aspartate to interact with R169 (Figure 7a, iv.). Thus, the carboxy-terminus of the retrieval signal sequentially engages R169, R5 and finally R47 (Figure 7c). Movement of the −4 position lysine towards E117 is concomitant with the final engagement of the carboxyl-terminus of the signal by R47, whereas D50 does not come in close proximity to the KDEL signal and there is only a transient interaction of S54 with the −5 position (Figure 7d).

**Figure 7.**
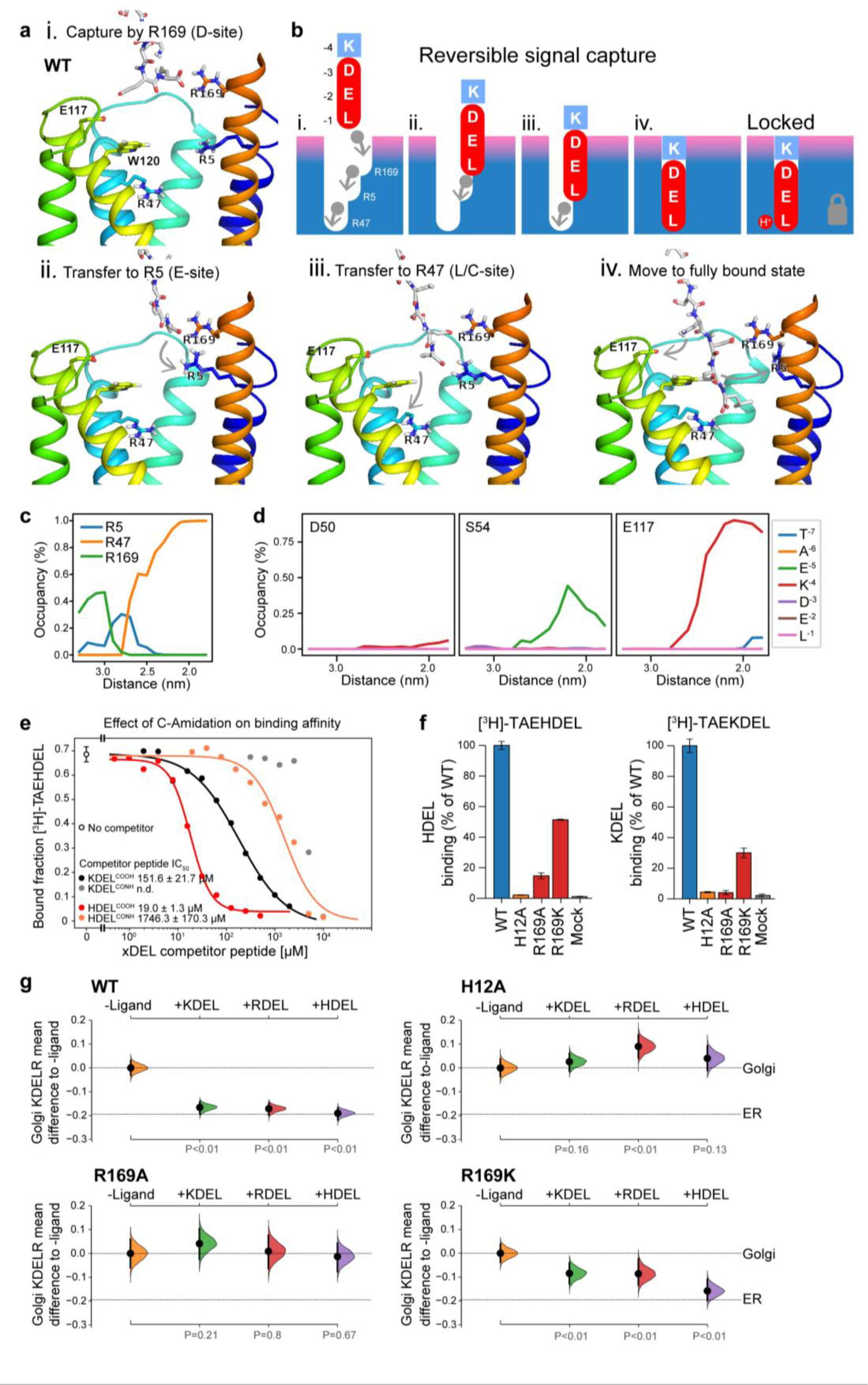
A baton-relay mechanism for initial retrieval signal capture by the KDEL receptor. **a.** Images depicting the key stages (i.-iv.) of KDEL binding to the wild-type (WT) human receptor simulated using molecular dynamics. Initial engagement of the C-terminus to R169 (i) is followed by transfer to R5 (ii), shortly followed by interaction of E −2 with R169 (iii). Finally, R47 engages the C-terminus allowing D −3 to interact with R169 (iv). **b.** A carton model depicting the key stages of retrieval signal binding and final pH-dependent locked state. **c.** Occupancy of the hydrogen bonds between the C-terminus of the KDEL retrieval signal and R5, R47 and R169 is plotted as a function of signal position within the binding pocket. **d.** The occupancy of potential hydrogen bonds between the different positions of the KDEL retrieval signal and D50, S54 and E117 is plotted as a function of signal position within the binding pocket. **e.** Competition binding assays for [^3^H]-TAEHDEL and unlabelled TAEKDEL and TAEHDEL with a free (COOH) or amidated (CONH) C-terminus to chicken KDELR2 showing IC_50_ values for the competing peptides. **f.** Normalised binding of [^3^H]-TAEHDEL and [^3^H]-TAEKDEL signals to the purified WT H12A, R169A or R169K mutant chicken KDELR2. A mock binding control with no receptor indicates the background signal. **g.** Distribution of WT, H12A, R169A and R169K KDEL receptors was measured in the absence (-ligand) or presence of K/R/HDEL^sec^. The mean differences for K/R/HDEL comparisons against the shared no ligand control are shown with sample sizes and p values.

We therefore propose a carboxyl-handover model for signal capture mediated by the ladder of arginine residues in the binding pocket (Figure 7b). As the carboxyl-terminus progresses further into the receptor binding site, the carboxylate groups at positions −2 and −3 engage their respective positions in the D- and E-sites respectively. Only the final stage of the binding, where the receptor closes around the signal locking it in place is pH dependent, all other stages are predicted to be freely and rapidly reversible. Because many proteins have a free C-terminal carboxylate, this highlights the importance of an initial proofreading stage where non-cognate signals are rejected, as we have already argued, due to their net charge.

To test these ideas, we investigated the importance of the retrieval signal C-terminus and R169 in the receptor using *in vitro* binding assays and functional experiments in cells. First, we synthesised C-terminally amidated HDEL and KDEL peptides and assayed their ability to bind to wild-type receptors (Figure 7e). Blocking the C-terminal carboxylate in this way completely abolished binding to KDEL and reduced the affinity for the HDEL peptide by two orders of magnitude from 19 ± 1.3 µM to 1.7 ± 0.1 mM. For HDEL, this residual affinity suggests the peptide still enters and exits the binding pocket, but fails to trigger the final pH dependent capture. Next, we performed binding assays with R169 variant receptors. Comparable results to the C-amidated peptide binding assays were obtained with R169A, which showed no binding to KDEL and greatly reduced binding to HDEL ligands (Figure 7f). Conservative substitution to R169K greatly reduced binding of both HDEL and KDEL in line with predictions (Figure 7f). Finally, we tested the R169 variants in ER retrieval assays. R169A mutant receptors showed no response to KDEL and only ∼10% response to HDEL signals (Figure 7g, Figure 7 – supplement 1a and b). By contrast, the conservative substitution R169K showed an attenuated response to both signals, in agreement with the simulation and reduced binding affinity (Figure 7g, Figure 7 – supplement 1a and b). We therefore conclude that the interaction of receptor R169 with the C-terminal carboxylate of the retrieval signal plays an important role in initial signal capture.

## DISCUSSION

### A baton-relay mechanism for initial signal capture by the KDEL receptor

Canonical ER retrieval signals can be broken down into two components: the −4 position, which enables the receptor to distinguish between different populations of ER proteins, and a tri-carboxylate moiety formed by the −3 aspartate, −2 glutamate and −1 C-terminal carboxylate. We propose a baton-relay handover mechanism for capture of this signal by the KDEL receptor wherein a ladder of three arginine residues in the receptor pairs with the three-carboxyl groups of the signal (Figure 8). During cargo capture, the receptor engages the retrieval signal in a stepwise process, with the C-terminal carboxyl group of the cargo protein moving between these three interaction sites. At neutral pH, C-terminal sequences will rapidly sample the binding site, a process that we imagine will occur in both the ER and Golgi apparatus. Only at acidic pH are cargo proteins captured once the C-terminus of the signal engages with R47 in the receptor, which then undergoes a conformational change to close around the signal, locking it in place, as we have explained previously (Bräuer et al., 2019). This mechanism explains why the retrieval signal must be located at the C-terminus of the cargo protein, and the defined requirement for either glutamate or aspartate residues at the −2 and −3 positions due to their carboxyl group containing side chains. Variation at the −4 position would not directly alter this initial capture mechanism, possibly explaining why it is the key determinant for signal selectivity.

**Figure 8.**
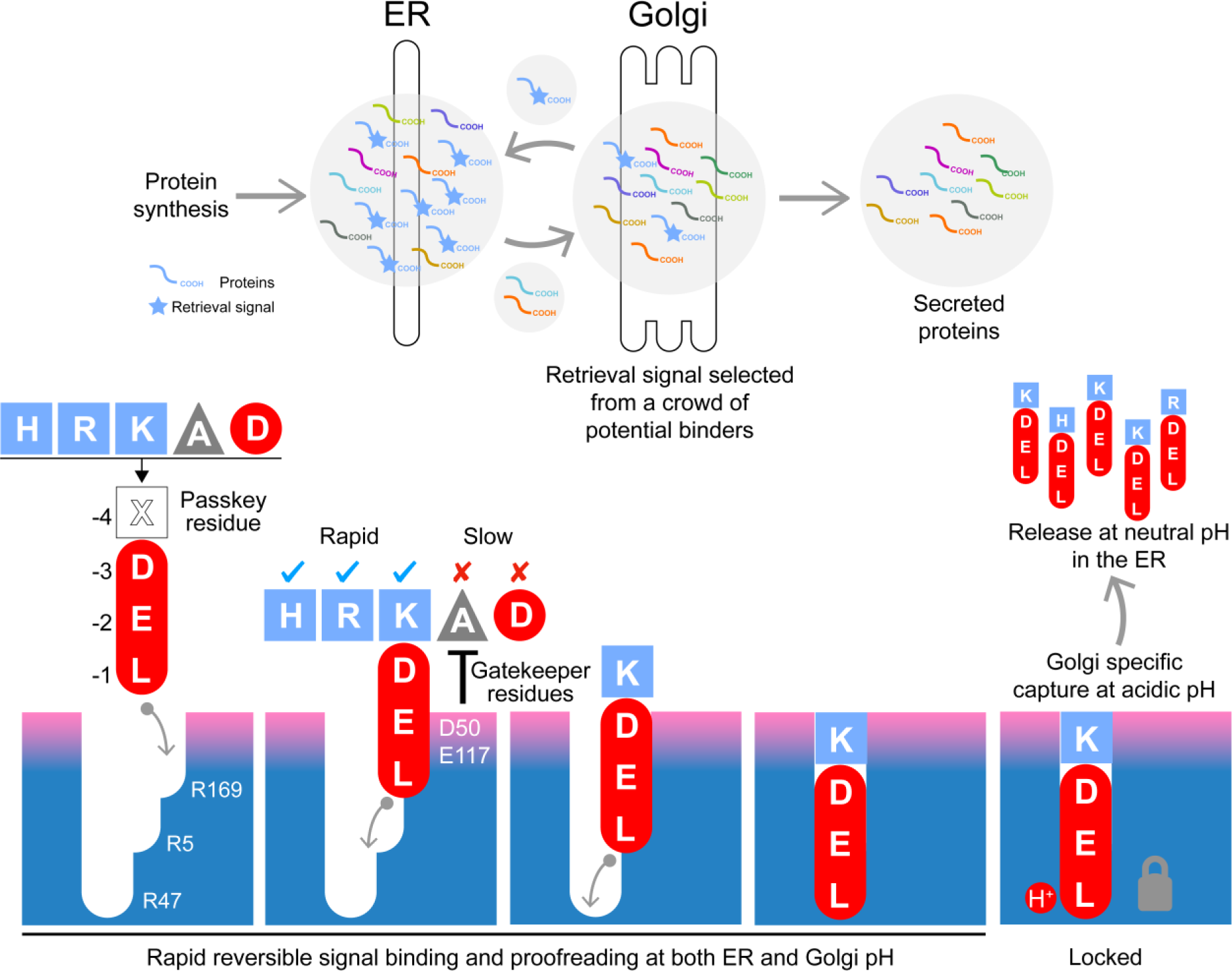
A combined proofreading and baton-relay handover model for initial signal capture by the KDEL receptor. Newly synthesized secretory and ER luminal proteins are translocated into the ER and on to the Golgi. Those proteins with C-terminal retrieval signals are captured by the KDELR receptor and returned to the ER. Other proteins with different C-terminal sequences move on to be secreted. The retrieval signal can be broken down into two sections: the variable −4 passkey position and the −1 to −3 positions with free carboxyl-terminus. Signals are initially captured through their free carboxyl-terminus by the receptor R169. This is then handed over to R5 and finally R47 in a baton-relay mechanism. Sequences are proofread for the residue at the −4 position by gatekeeper residues D50 and E117. Unwanted signal variants are rejected. Only signals that completely enter the binding pocket and engage R47 can undergo pH dependent capture and return to the ER.

The structures we have obtained for the KDEL receptor with bound HDEL, RDEL or KDEL signals reveal that the side chains at the −4 position form a salt bridge interaction with E117 but, crucially, not D50 as previously proposed. Unexpectedly, the salt bridge interaction between E117 and the −4 position of the retrieval signal makes only a limited contribution to binding affinity and does not explain the higher affinity for HDEL. Our other data show that the higher affinity for HDEL is due to the stronger π-π interaction between the histidine of the −4 position of the retrieval signal and W120 in the receptor. However, because E117A mutant receptors have expanded specificity and can recognise ADEL, we conclude that the side chain at the −4 position is unlikely to play a major role in binding affinity for signals other than HDEL. For these reasons, we refer to the −4 position as the passkey residue, important for the signal selection only. By determining net charge on the signal it may thus play a greater role in initial binding kinetics, rather than affinity per se.

Previous work has suggested that the −5 and −6 positions of the retrieval signal also play a key role in signal binding (Alanen et al., 2011), and that the individual human KDEL receptors have unique specificity (Raykhel et al., 2007). However, these properties are not completely consistent with the structures, pattern of sequence conservation, or wider analysis presented here. It is noteworthy that both those studies used a bimolecular fluorescence complementation approach where the signal and receptor are dimerised by a split YFP molecule that will likely contribute to the observed signal binding affinity. This will interfere with the initial proofreading mechanism described here, making comparison with our data difficult. Since the three human KDEL receptors share identical ligand binding residues and only differ in conservative substitutions at position 54, we believe they will have the same or closely-related ligand binding properties. Based on the structures it seems reasonable that the −5 position may contribute to signal proofreading in some cases. However, as we show, a wide variety of signals that lack any obvious conserved features upstream of the canonical tetrapeptide function efficiently to trigger ER retrieval of the receptor (Figure 1a and S5b-S5e), suggesting the −5 position modulates but does not play an essential role in signal recognition. Taken together, these data support a model for retrieval sequence recognition that explains both the importance of the free C-terminal carboxyl group and how changes at the −4 position can modulate binding to the receptor.

### The important role of luminal pH in regulating HDEL-mediated ER retrieval

One important outcome from our work is the idea that KDEL receptors are not optimised for an individual signal and must retain the ability to differentiate variant high and low affinity ER retrieval signals. We propose that cells exploit these properties to maximise the retrieval efficiency of a broad range of ER resident proteins with widely different abundance, over 2 or 3 orders of magnitude. This idea explains an explanation for the functional significance of the affinity differences of retrieval signal variants in mammalian cells. The most abundant proteins use the KDEL retrieval signal, whereas lower abundance proteins tend to carry the HDEL signal. By artificially increasing the concentration of HDEL proteins we can show that this effectively poisons the ER retrieval system, leading to the secretion of normally retained ER chaperones. This behaviour is reminiscent of other cellular regulatory systems, where substrate or signal binding properties are optimised for rate and turnover, rather than for the highest affinity which can reduce throughput of the pathway. Indeed, in some cases electrostatic properties are exploited to create rapid-binding high-affinity inhibitors that outcompete substrates (Cundell et al., 2016; Schreiber and Fersht, 1996). This may explain why histidine has been selected for the highest affinity variant of the signal to counteract this effect. For the HDEL variant of the retrieval signal, protonation of both the receptor histidine 12 and signal peptide histidine −4 favour binding to the receptor in the Golgi. However, deprotonation of both the retrieval signal and receptor at pH 7.0 enable rapid release in the ER, and hence receptor recycling to the Golgi. Thus, HDEL binds more tightly than KDEL in the Golgi, but still releases rapidly in the ER. A signal with the same affinity as HDEL that was not proton dependent would strongly inhibit retrieval even at low concentration due to slow release at neutral pH. An alternative mechanism to capture low abundance ER proteins would have been to increase the cellular concentration of the KDEL receptor from the observed low levels. That would require receptors to be nearly stoichiometric with cargo, a problematic proposition considering the millimolar concentration of ER chaperones. These potential traps are avoided by the combination of pH-regulation of both the receptor and the high affinity HDEL signal. Thus, the versatile binding site architecture of a single KDEL receptor enables differentiation of both high and low affinity signals, thereby enabling efficient ER retrieval of both low and high abundance proteins in eukaryotic cells.

## ACKNOWLEDGEMENTS

We thank the staff at I24 Diamond Light Source, UK for access to the beamline. This work was supported by Wellcome awards (219531/Z/19/Z, 203741/Z/16/A and 109133/Z/15/A) to SN, FAB and PCB. Computation time was provided by JADE (EP/P020275/1) and ARCHER via HECBioSim (http://www.hecbiosim.ac.uk), supported by an EPSRC grant (EP/R029407/1) to PCB. ZW is a Wellcome Trust PhD student (203741/Z/16/A)

The authors declare no competing financial interests.

## AUTHOR CONTRIBUTIONS

FAB, SN, PB, AG and JLP designed the experiments. PB, AG, TS, JLP, FAB and SN carried out experiments and interpreted the data. ZW and PCB designed and performed the computational analysis. FAB and SN wrote the paper with input from all authors.

## MATERIALS & METHODS

### RESOURCES TABLE

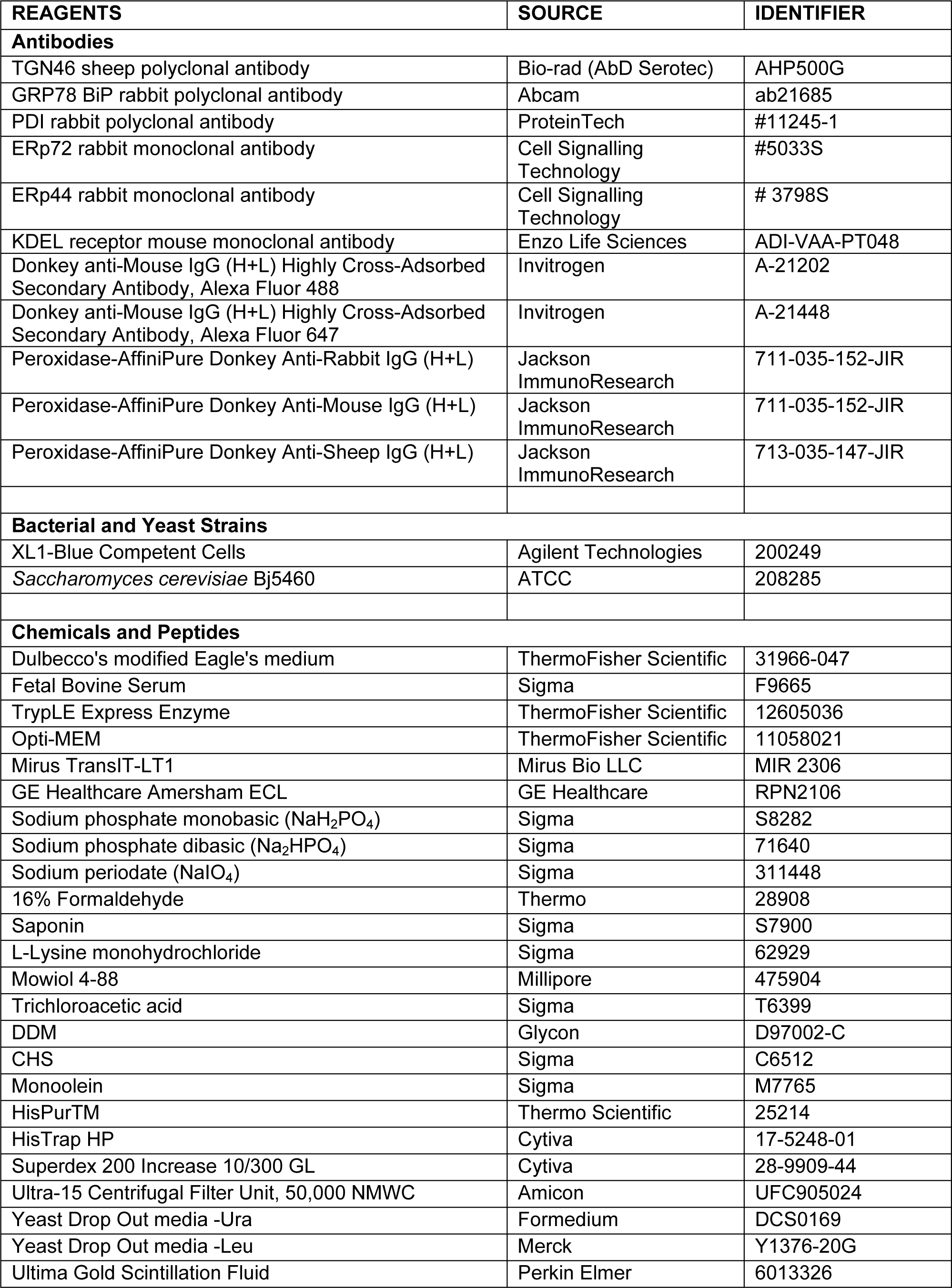

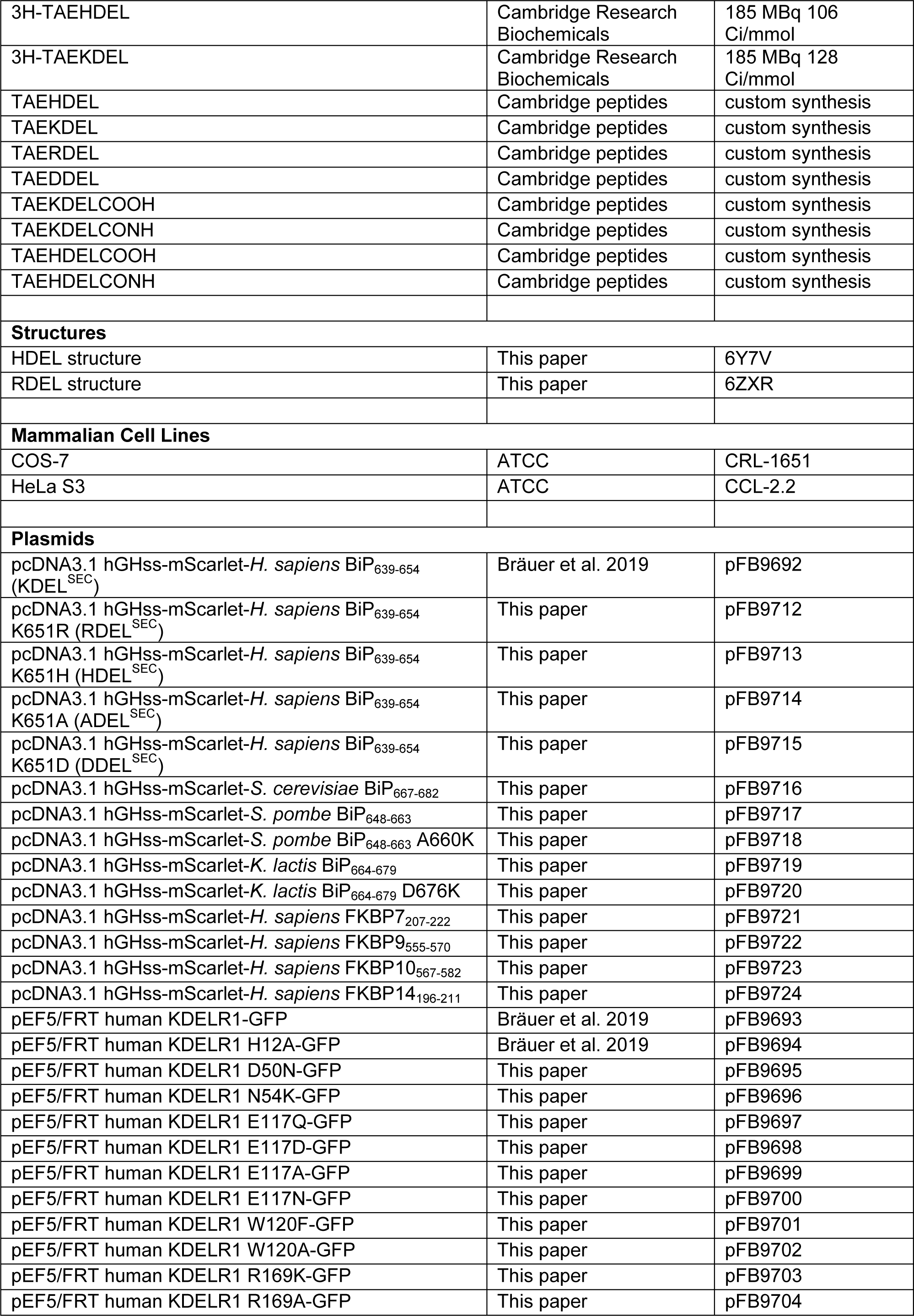

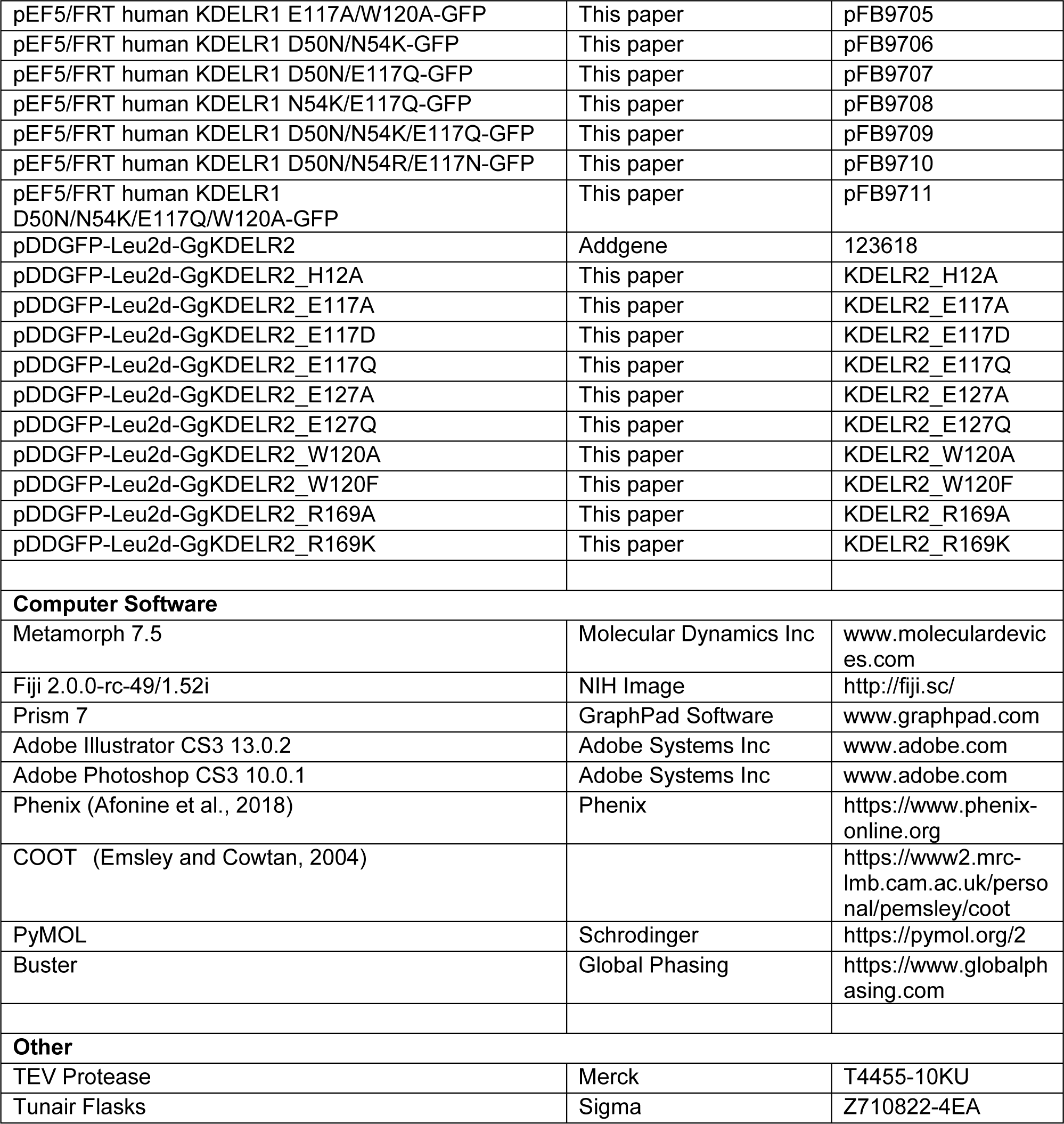

#### Mammalian cell lines

HeLa cells were cultured at 37°C and 5% CO_2_ in DMEM containing 10% [vol/vol] foetal bovine serum (Invitrogen). For passaging, cells were washed in PBS, and then removed from the dish by incubation with TripLE Express (Thermo Fisher Scientific).

#### ER retrieval and secretion assays

*Homo sapiens* KDELR1 (Uniprot: P24390) was cloned into the pEF5/FRT low level mammalian expression vector with a C-terminal 20 amino acid linker made up of 5 copies of Gly-Ser-Ser-Ser followed by GFP to create KDELR-GFP. Specific point mutations in the ligand binding site, described in the figures, were introduced using the Quickchange protocol (Stratagene). To create the mScarlet-KDEL^sec^ ligand construct, mScarlet with the N-terminal hGH signal peptide and the 16 C-terminal residues of human BiP at its C-terminus, containing the KDEL signal, was cloned into the pcDNA3.1 vector. This was modified using site-directed mutagenesis or annealed oligo ligation to create C-terminal retrieval signal variants from known human and yeast ER proteins. COS-7 cells were grown on 10 mm diameter 0.16-0.19 mm thick glass coverslips in DMEM containing 10% [vol/vol] bovine calf serum at 37°C and 5% CO_2_. Cells were plated at 50,000 cells per well of a 6-well plate, each well containing 2 coverslips. For ER retrieval assays, the cells were transfected after 24 h with 0.25 µg KDELR-GFP and 0.5 µg mScarlet-ligand (+ xDEL ligand) or 0.25 µg KDELR-GFP and 0.5 µg pcDNA3.1 (− ligand) diluted in 100µl Optimem and 3 µl Mirus LT1 (Mirus Bio LLC). After a further 18 h, cells were washed twice with 2 mL of PBS, then fixed for 2 h in 2 mL 2% wt/vol) formaldehyde in 87.5 mM lysine, 87.5 mM sodium phosphate pH 7.4, and 10 mM sodium periodate. Subsequently, coverslips were washed three times in 2 mL permeabilization solution 100 mM sodium phosphate pH 7.4, then permeabilised in 1 mg mL^-1^ BSA, 0.12 mg mL^-1^ saponin, and 100 mM sodium phosphate pH 7.4 for 30 min. Primary and secondary antibody staining was performed for 60 min in permeabilization solution at 22°C. Commercially available antibodies were used to detect the Golgi protein TGN46 (sheep; AbD Serotec). Coverslips were mounted on glass slides in Mowiol 4-88 and imaged with a 60×/1.35 NA oil immersion objective on an Olympus BX61 upright microscope (with filtersets for DAPI, GFP/Alexa-488, −555, −568, and −647 (Chroma Technology Corp.), a 2048×2048 pixel CMOS camera (PrimΣ; Photometrics), and MetaMorph 7.5 imaging software (Molecular Dynamics Inc.). Illumination was provided by a wLS LED illumination unit (QImaging). Image stacks of 3-5 planes with 0.3 µm spacing through the ER and Golgi were taken. The image stacks were then maximum intensity projected and the selected channels merged to create 24-bit RGB TIFF files in MetaMorph. To produce the figures, images in 24-bit RGB format were cropped in Photoshop to show individual cells and then placed into Illustrator (Adobe Systems Inc.). To determine ER retrieval efficiency, the Golgi signal (integrated pixel intensity) for the KDEL receptor was measured (Schindelin et al., 2012) in the region defined by the Golgi marker antibody in the presence (+) and absence (-) of ligand. Golgi signal was normalised to the ER signal, to account for different expression levels.

For ER secretion assays, upon transfection HeLa S3 cells were allowed to express the proteins for 24 h. The media were TCA precipitated and both cell and media were resuspended and boiled in SDS-PAGE sample buffer. All samples were analysed by Western blotting for xDEL ligand, resident ER chaperones BIP (rabbit #ab21685, Abcam), PDI (rabbit #11245-1, ProteinTech), ERP72 (rabbit #5033S, Cell Signalling Technology), ERP44 (rabbit #3798S, Cell Signalling Technology) and the KDEL receptor (mouse ADI-VAA-PT048, Enzo Life Sciences).

#### Statistical analysis of ER retrieval and secretion

To estimate the effect sizes and significance of receptor mutations for ligand-mediated ER retrieval, data was analysed in R using the open-source package dabestr (Ho et al., 2019; Team, 2017; Wickham, 2010). Data are presented as Cumming estimation plots, where the raw data is plotted on the upper axes and mean differences are plotted as bootstrap sampling distributions on the lower axes for 5000 bootstrap samples. Each mean difference is depicted as a dot. Each 95% confidence interval is indicated by the ends of the vertical error bars; the confidence interval is bias-corrected and accelerated. The p values reported are the likelihood of observing the effect size, if the null hypothesis of zero difference is true. For each permutation p value, 5000 reshuffles of the control and test labels were performed.

#### KDEL receptor crystallisation and structure determination

*G*g KDELR2 was expressed and purified as described previously (Bräuer et al., 2019), concentrated to 14.5 mg mL^-1^ and incubated with 6.4 mM TAEHDEL peptide on ice for one hour prior to crystallisation. Crystals were set up at 20 °C as above using precipitant 30% (v/v) PEG 600, 100 mM MES pH 6.0, 100 mM Sodium Nitrate. Phases were determined via molecular replacement using Phaser and employing PDB:6I6H as the search model with the TAEKDEL peptide removed from the search model. The TAEHDEL peptide was built into difference density using Coot (Emsley et al., 2010), followed by refinement in BUSTER (Blanc et al., 2004).

#### Retrieval signal binding assays

Binding assays were performed in 20 mM MES pH 5.4, 40 mM Sodium Chloride, 0.01% DDM 0.0005% CHS unless stated otherwise. 5 µL of ^3^H-TAEK/HDEL (Cambridge peptides, UK) at 20 nM was incubated with 5 µL of *Gg* KDELR or variants thereof at the desired concentration at 20 °C for 10 min. The reaction was then filtered through a 0.22 µm mixed cellulose ester filters (Millipore, USA) using a vacuum manifold. Filters were then washed with 2 x 500 µL buffer. The amount of peptide remaining bound was measured using scintillation counting in Ultima Gold (Perkin Elmer). Experiments were performed a minimum of three times to generate an overall mean and standard deviation. Data was normalised to the maximal binding at pH 5.4 and fit with a four-parameter logistic non-linear regression model.

#### Relative binding free energy calculations

To compute the free energy of the deprotonation of the histidine or lysine and the mutation of lysine to histidine, molecular mechanics based alchemical transformation was performed. The free energy difference was taken as the difference in the free energy of the transformation between the protein-peptide complex and the peptide in solution. The KDEL receptor in the protein-peptide complex was taken from the crystal structure (KDEL: 6I6B (Bräuer et al., 2019); HDEL: 6Y7V). The C-terminus of the receptor was modelled to full length using Modeller 9.21 (Webb and Sali, 2016); 100 models were created and the one with the best DOPE score was selected (Shen and Sali, 2006). The protein was then embedded into a lipid membrane containing 186 DMPC lipids using the procedure described by us previously (Wu et al., 2019). The system of peptide in solution was constructed by taking the coordinates of the peptide from the crystal structure and placing in a box, where the box edge was at least 2 nm from the peptide. Both systems were solvated and neutralised to final salt concentration of 150 mM NaCl. For the deprotonation calculations, the change in charge in the system was counteracted by simultaneously charging a sodium ion in the corner of the box (ie at the start of the process the charge was zero and by the end it was +1). To minimise the interactions between the histidine (or lysine) and this alchemical sodium ion, the histidine/lysine residues were restrained to the centre of the box via their Cα atom using a harmonic restraint of 1000 kj/mol/nm and the alchemical sodium ion was either restrained to the edge of the box for the peptide in solution or restrained to the z-axis in the case of the peptide-protein complex.

The Amber ff14SB force field (Maier et al., 2015) was used to describe the protein and alchemical transformation was done with pmx (Gapsys et al., 2015). Lipids were described by LIPID17, which was ported from amber to Gromacs by us (Wu and Biggin (2020). GMX_lipid17.ff: Gromacs Port of the amber LIPID17 force field. Zenodo. http://doi.org/10.5281/zenodo.3610470). The simulations were run with GROMACS 2018 (Abraham et al., 2015). The simulation input parameters were set according to recommendations suggested by pmx. Since the equilibrium method was used, the sc-alpha and sc-sigma parameters were set to 0.5 and 0.3 respectively. For the lysine to histidine transformation, a total of 21 lambda windows with 0.05 equal spacing were used to transform the charge and the vdw parameters at the same time. A soft-core potential was used for the coulombic interactions to avoid singularity effects. For the deprotonation calculations, 11 equally spaced windows were used to change the partial charge and an addition window was used to complete the transformation. After energy minimisation, each window was run for 200 ps in the NVT ensemble and 1 ns in an NPT ensemble with position restraints of 1000 kj/mol to reach a final temperature of 310 K and 1 bar. 30 ns production runs with replicate exchanges at intervals of 1 ps were then performed. Data were analysed using the Multistate Bennett Acceptance Ratio with alchemical analysis with the first 5 ns discarded (Klimovich et al., 2015). For each transformation, three replicates were performed and the result was given as the mean and standard deviation. For the LYS/HIP transformation, since both HDEL-bound and KDEL-bound structure were available, six simulations (three starting from KDEL-bound structure and three from HDEL-bound structure) were used to produce the results.

To compute the free energy difference of KDEL to HDEL transformation, the total free energy difference of alchemically changing KDEL to HDEL is computed as

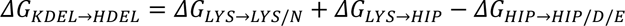

Where 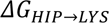 is the free energy difference of converting KDEL to HDEL when both lysine and histidine are in the protonated form. 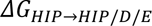 and 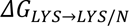 is the free energy of converting protonated histidine or lysine from the protonated to an ensemble of protonated and deprotonated forms (for example we might calculate the energy to go from 100% protonated to an ensemble of 40% protonated and 60% deprotonated):

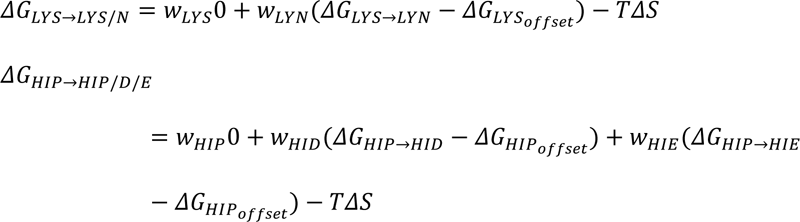

The 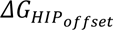 and 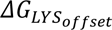 are terms to calibrate the computational protonation free energy to the experimental microscopic pka (histidine: 6.0; lysine: 8.95) and were defined as:

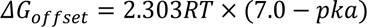

*w* is the Boltzmann weight of each protonation state and is computed as:

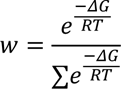

and *ΔS* is the configurational entropy and is defined as:

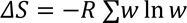

#### Quantum mechanical calculations for the effect of HDEL protonation

To explore the interactions between the signal and receptor, the histidine of the HDEL signal and tyrosine (W120) of the receptor were taken from the crystal structure and capped at both ends (with acetyl and amide groups the N and C-termini respectively). The hydrogens were added to the complex and the three different protonation sates of the histidine were constructed with Maestro 2019.2. The capped three amino acid complex were geometry minimised with non-hydrogen atoms constrained at the RI-B3LYP-D3(BJ)/def2-TZVP theory level with geometry counterpoise (Grimme et al., 2010; Grimme et al., 2011; Kruse and Grimme, 2012; Weigend, 2006; Weigend and Ahlrichs, 2005) using ORCA 4.2.0 (Neese, 2012). The interactions between the three different protonation states of the histidine and W120 were computed at the SAPT2+/jun-cc-pVDZ (Parker et al., 2014) theory level from the geometry optimised structure using psi4 1.3.2 (Parrish et al., 2017).

#### Simulation of signal engagement with the binding site

To obtain a converged view of how the KDEL peptide enters the KDEL receptor, umbrella sampling was used to enhance the sampling of the behaviour of the C-terminus in the binding pocket. The initial frames were generated by pulling the N-terminus of the KDEL peptide out of the binding pocket using a moving restraint (Gromacs 2019.4/plumed 2.6.0) (consortium, 2019). The collective variable (CV) was defined as the distance between the N-terminus of the KDEL peptide (N atom) and the centre of the binding pocket, which was defined as the centre of the Cα atoms of residue 9, 44, 64, 124 and 162. Pulling was performed using a CV=1.8 nm to 3.3 nm with a restraint strength of 1000 kcal/mol/nm for 100 ns. To prevent the complete dissociation of the peptide from the receptor, a one-side distance restraint was applied on the distance between the C-terminus of the peptide (atom C) and the binding pocket at 1.7 nm with a strength of 1000 kj/mol/nm. Sixteen windows were set up where the CV was varied from 1.8 nm to 3.3 nm with a step of 0.1 nm and were run for 500 ns. The results were analysed with MDAnalysis 1.0 (https://conference.scipy.org/proceedings/scipy2016/oliver_beckstein.html).

#### Quantification and statistical analysis

Details of the number of experimental repeats, numbers of cells analysed and the relevant statistics are detailed in the figure legends and specific method details.

## FIGURE SUPPLEMENTS

**Figure 1 – supplement 1.**
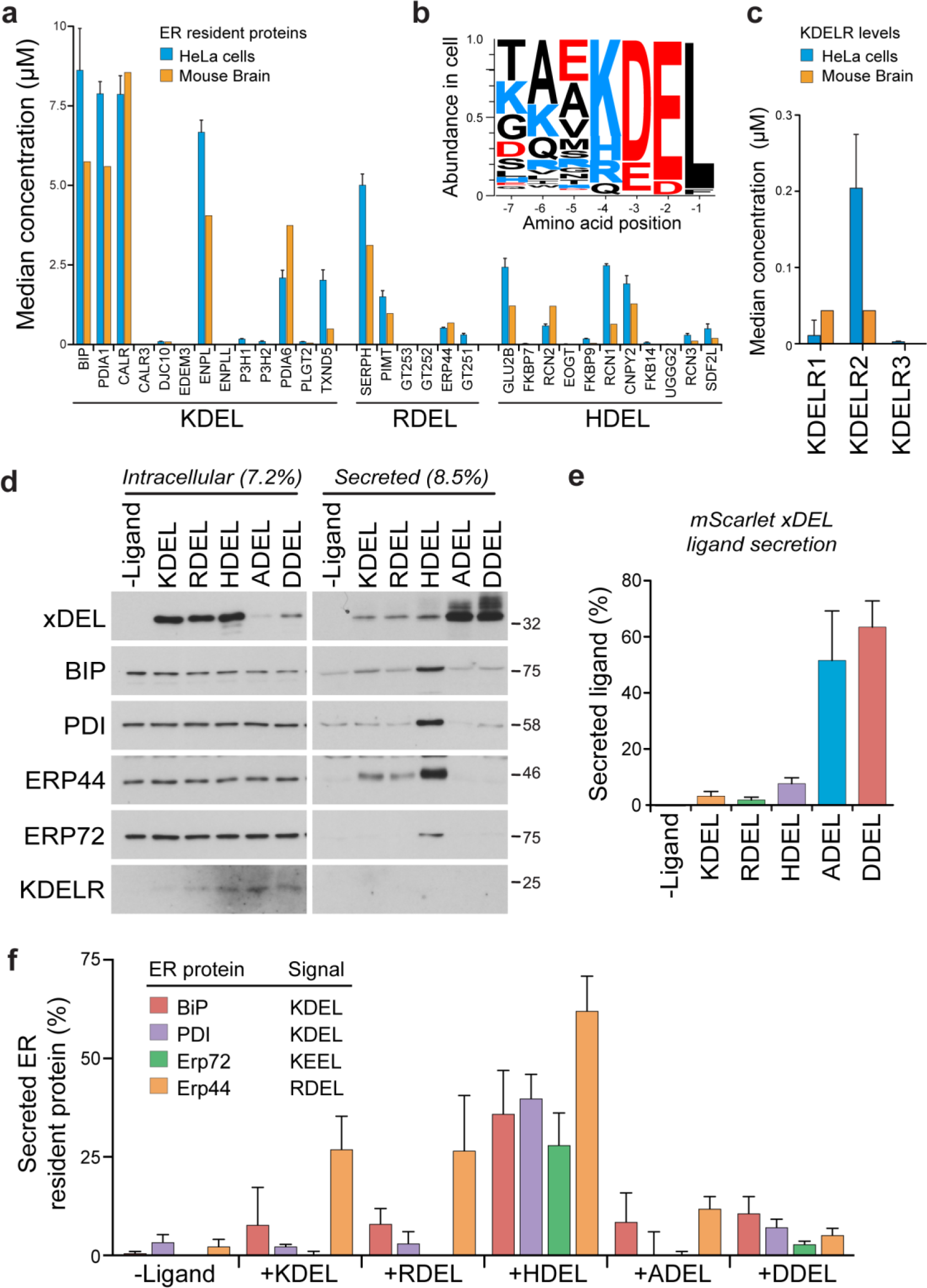
Abundance of ER resident proteins and chaperones in human cells and mouse brain. **a.** The mean concentration of ER resident chaperones with the indicated ER retrieval sequence variant is plotted in the bar graph (Itzhak et al., 2017; Itzhak et al., 2016). **b.** Combined cellular concentrations of ER resident proteins with canonical KDEL, RDEL and HDEL retrieval sequences in HeLa cells and mouse brain. **c.** The mean concentration of KDELR1, KDELR2 and KDELR3 in HeLa cells and mouse brain is plotted in the bar graph (Itzhak et al., 2017; Itzhak et al., 2016). **d.** Cells and media collected from cultures expressing the xDEL variants (mScarlet-xDEL^sec^) indicated in the figure were Western blotted for resident ER chaperones and KDELR. **e.** A bar graph of xDEL secretion showing mean ± SEM (n=3). **f.** Endogenous ER chaperone secretion was measured by western blotting after challenge with different retrieval signals, and plotted as a bar graph showing mean ± SEM (n=3).

**Figure 2 – supplement 1.**
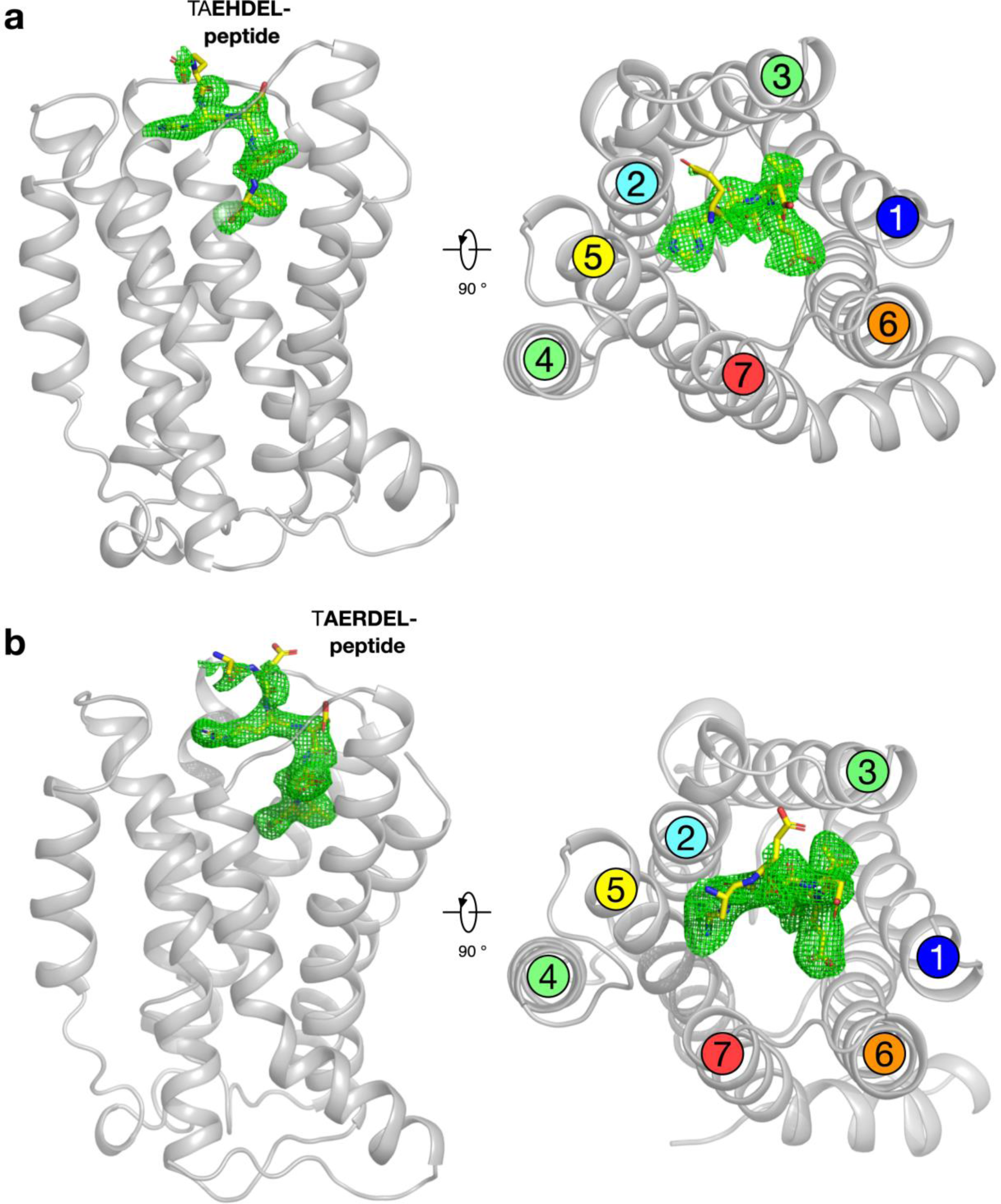
Polder difference density electron density maps for HDEL and RDEL peptides. **a.** The structure of the KDELR bound to the TAEHDEL peptide is shown as in Figure 2a. The *m*Fo-*D*Fc difference electron density used for model building is displayed (green mesh), contoured at 3σ. **b.** Equivalent maps calculated for the RDEL peptide.

**Figure 3 – supplement 1.**
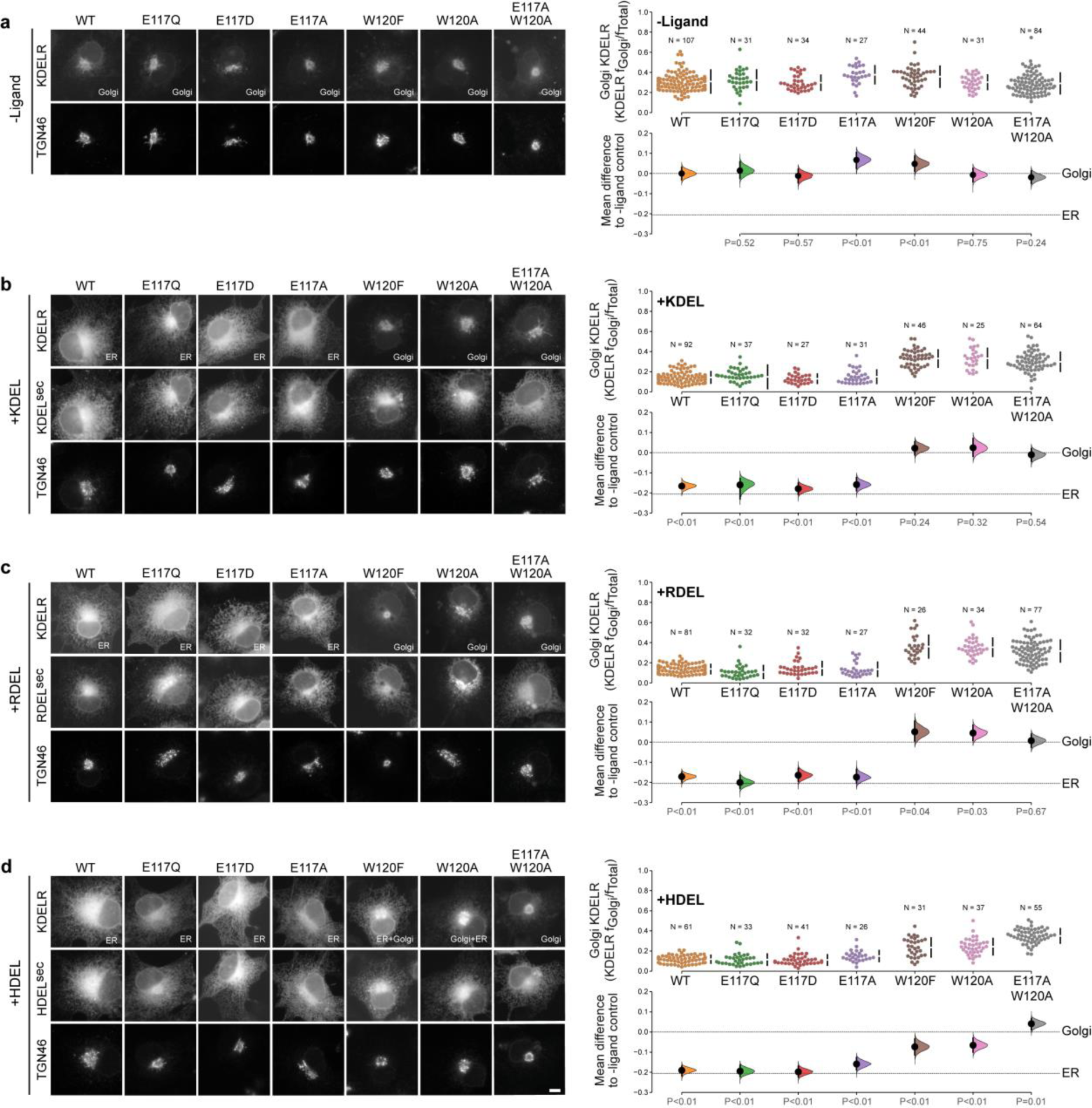
Effect of KDEL receptor E117 and W120 mutants on retrieval signal function in cells. **a.** Distribution of WT, E117 and W120 mutant KDEL receptors was measured in the absence (-ligand) or presence of **b.** KDEL, **c.** RDEL or **d.** HDEL retrieval signals (K/R/HDEL^sec^). TGN46 was used as a Golgi marker. Scale bar is 10µm. The fraction of WT, E117 and W120 mutant KDEL receptor localised to the Golgi was measured before (no ligand) and after challenge with different retrieval signals (K/R/HDEL) as indicated. Golgi signal for KDEL receptor and ligand intensity are shown on the scatter plots with sample sizes. Effect sizes are shown as the mean difference for K/R/HDEL comparisons against the shared -ligand control with sample sizes and p-values. The Cumming estimation plots for this data are used in main Figure 3c.

**Figure 4 – supplement 1.**
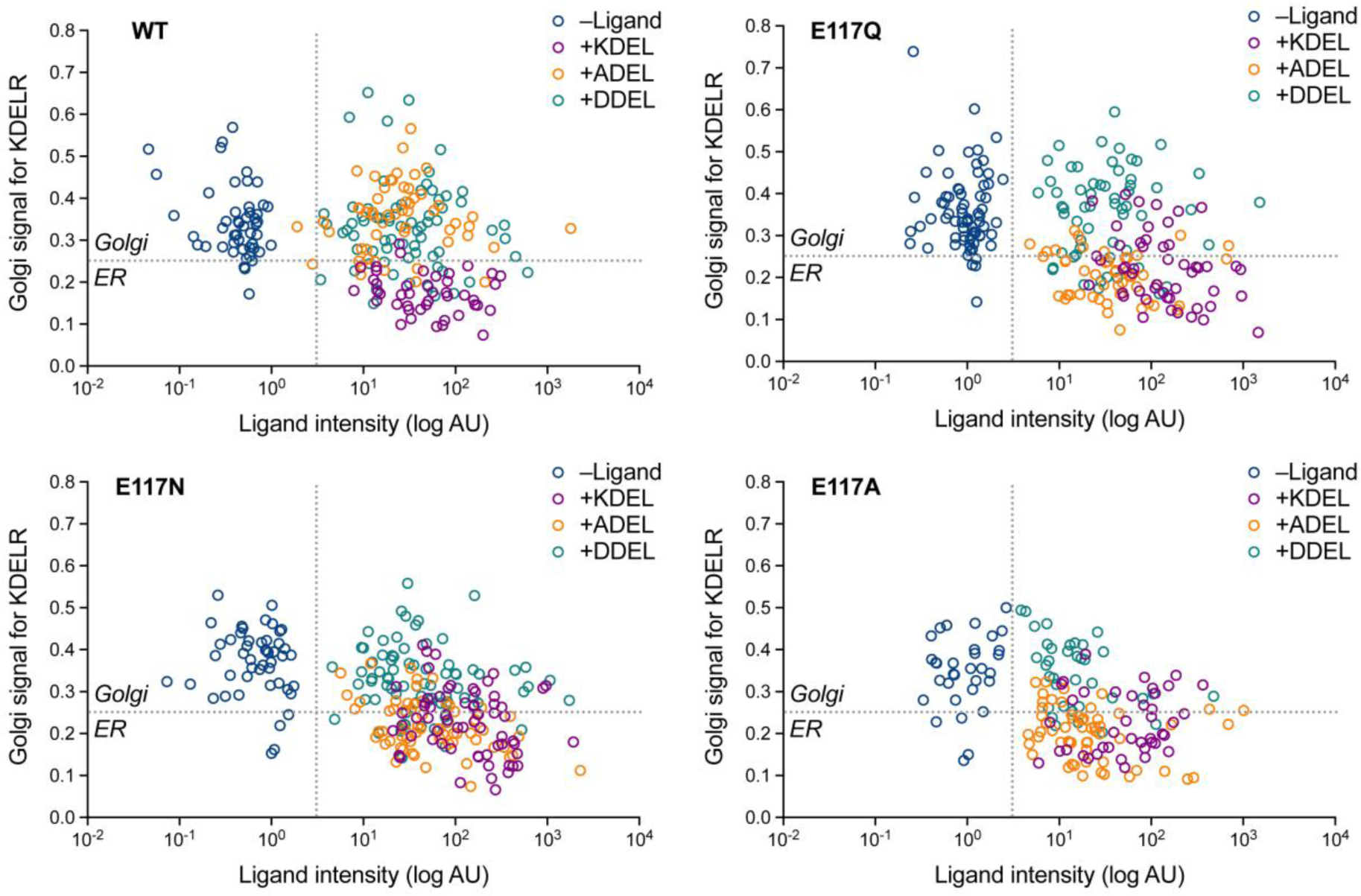
Effect of ligand levels on the response of KDEL receptor E117 mutants to KDEL, ADEL and DDEL signals. Distribution of WT, E117A, E117Q and E117N mutant KDEL receptors was measured in the absence (-ligand) or presence of KDEL, ADEL and DDEL retrieval signals. The fraction of WT, E117 mutant KDEL receptor localised to the Golgi was measured before (no ligand) and after challenge with different retrieval signals. Ligand intensity was also measured and plotted against the Golgi fraction of KDEL receptor.

**Figure 5 – supplement 1.**
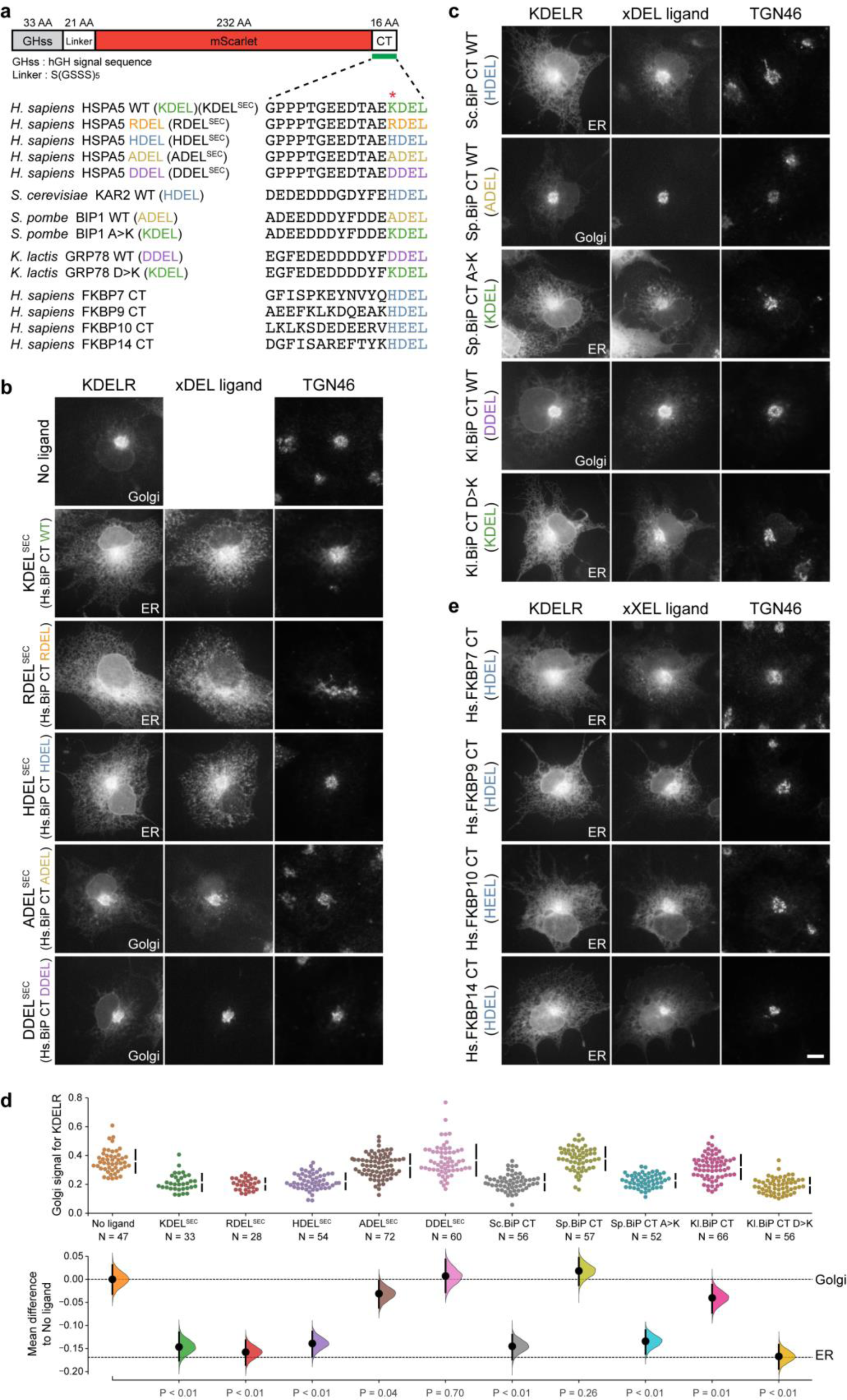
Comparison of human and yeast ER retrieval signals. **a.** Schematic of the ER retrieval construct showing the human growth hormone signal sequence (hGHss), linker, mScarlet fluorescent protein, and 16 amino acid extension carrying a C-terminal (CT) retrieval signal from known human and yeast ER proteins. A sequence alignment shows the conservation of the retrieval signal. **b.** WT KDEL receptor distribution was followed in the absence (-ligand) or presence of human BIP derived signals (K/R/H/A/DDEL^sec^). **c.** WT KDEL receptor distribution was followed in the absence (-ligand) or presence of yeast BIP derived signals K/R/HA/DDEL^sec^. **d.** The fraction of KDEL receptor localised to the Golgi was measured before (no ligand) and after challenge with the different retrieval signals tested in b. and c. Effect sizes are shown as the mean difference for retrieval signal comparisons against the shared -ligand control with sample sizes and p-values. **e.** WT KDEL receptor distribution was followed in the absence (-ligand) or presence of ER retrieval signals from human FKBP family proteins. In all image panels, TGN46 was used as a Golgi marker and the scale bar is 10µm.

**Figure 6 – supplement 1.**
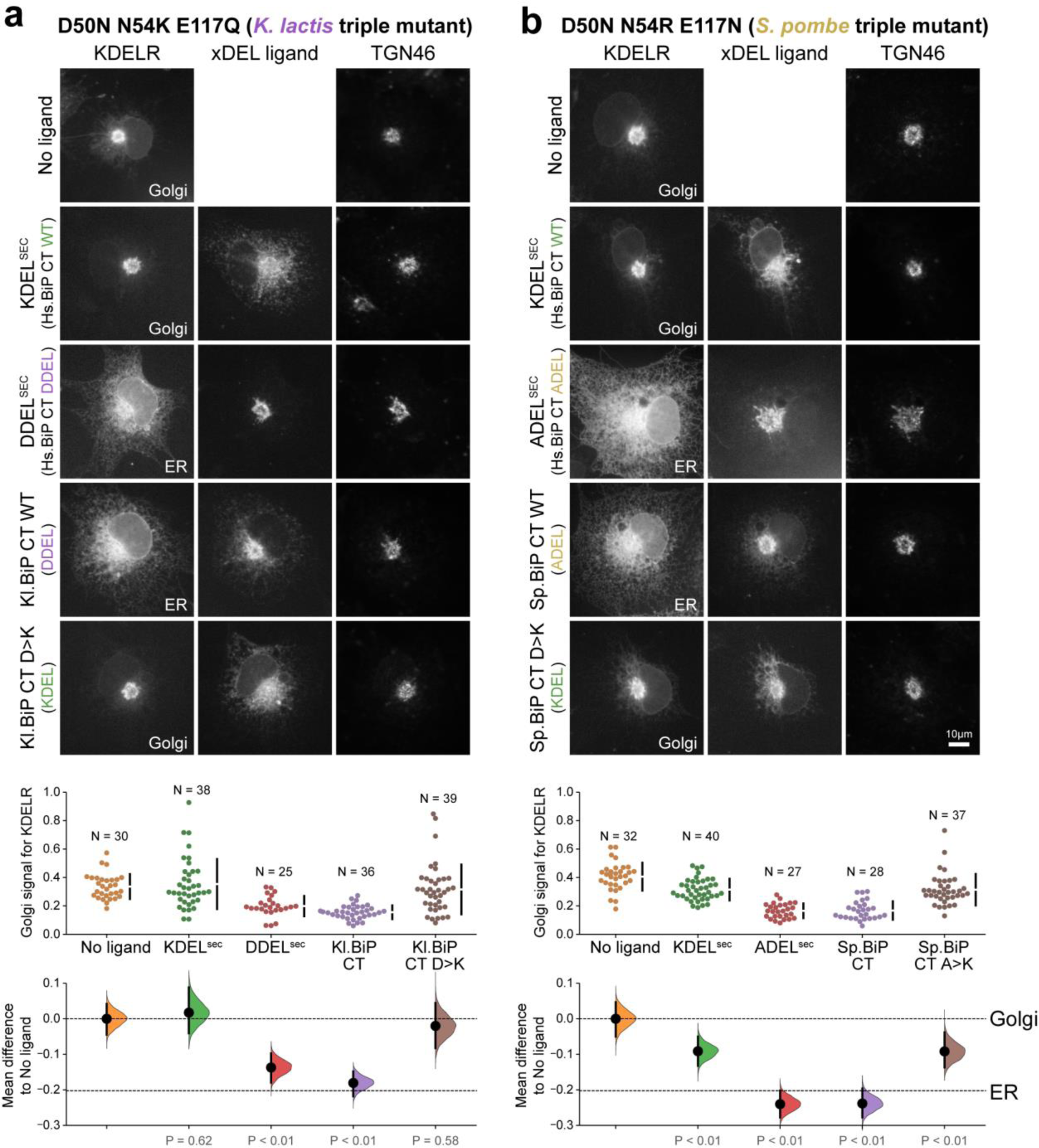
Retrieval specificity of “*K. lactis*” and “*S. pombe*” triple mutant KDEL receptors. **a.** Triple mutant “*K. lactis*” and **b.** “*S. pombe*”-like KDEL receptors were tested for K/A/DDEL-induced redistribution from Golgi to ER. KDEL receptor distribution was followed in the absence (-ligand) or presence of the indicated ligands. TGN46 was used as a Golgi marker. Scale bar is 10µm. The fraction of WT and mutant KDEL receptor localised to the Golgi was measured before (no ligand) after challenge with different retrieval signals.

**Figure 7 – supplement 1.**
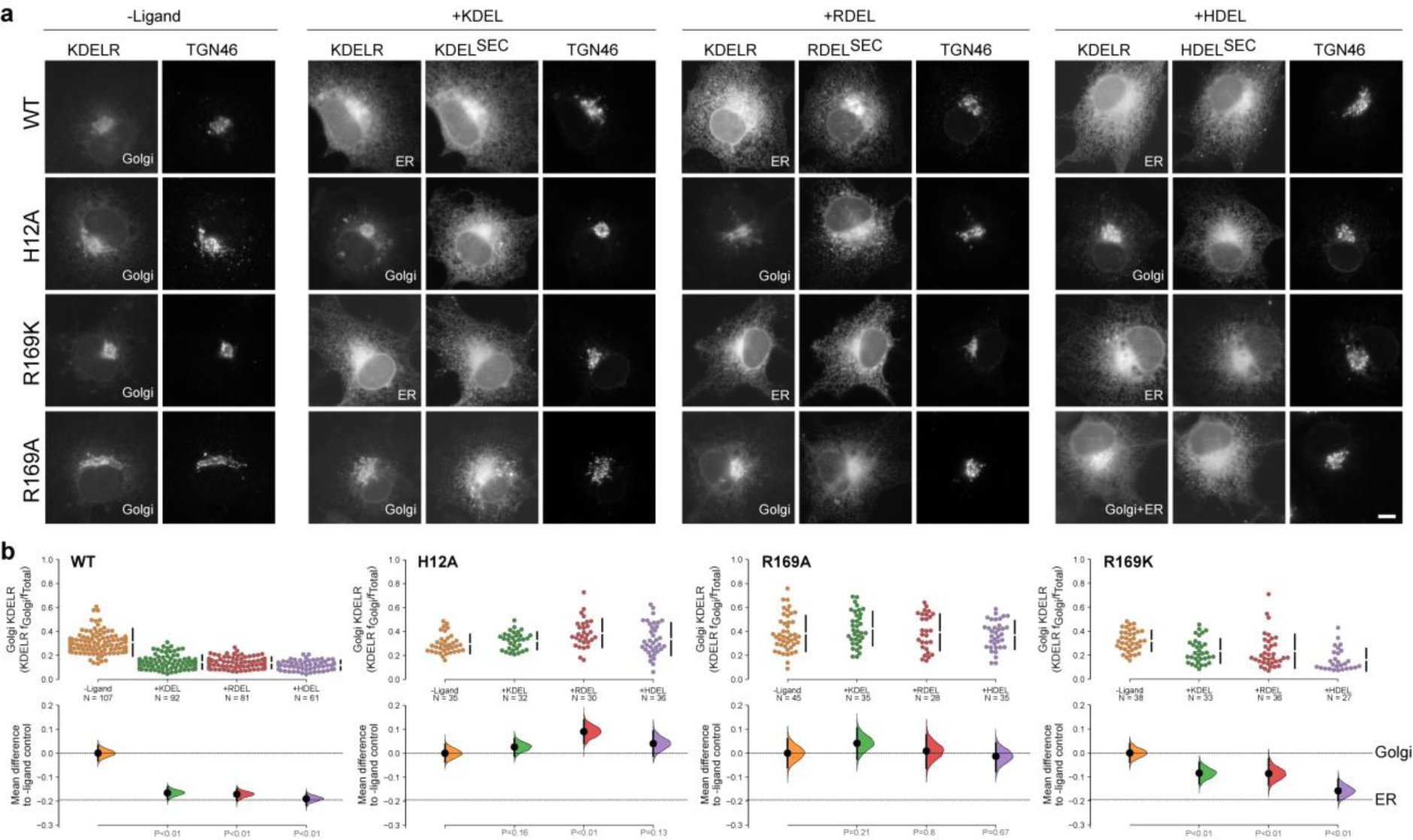
R169 plays a crucial role in signal recognition. **a.** Distribution of WT, H12A, R169A and R169K KDEL receptors was measured in the absence (-ligand) or presence of K/R/HDEL (mScarlet-xDEL^sec^). TGN46 was used as a Golgi marker. Scale bar is 10µm. **b.** The raw data for the fraction of KDEL receptor fluorescence in the Golgi is plotted on the upper axes with sample sizes, with effect sizes shown in the lower graphs.

